# Targeting CDK12 condensates at nuclear speckles with rationally designed cyclin K molecular glue degraders

**DOI:** 10.1101/2025.03.31.646293

**Authors:** Kaiyan Ye, Qinyang He, Yan Cheng, Meiying Zhang, Xiaonuo Ma, Huiling Mu, Mei-Chun Cai, Yuqiao Hou, Kaijie Zhou, Huaijiang Xiang, Shaoqing Zhou, Yunxia Li, Yinuo Zang, Jingxin Lv, Lin Cheng, Zhiliang Wang, Juan Liao, Xi Wang, Dayong Chen, Bing Yang, Fan Yang, Libing Xiang, Dongxin Zhao, Zhen Wang, Ke Ding, Yaoyang Zhang, Ying Li, Xia Yin, Wen Di, Yixiao Zhang, Peiye Shen, Guanglei Zhuang, Li Tan

## Abstract

Transcriptional machinery often operates in the configuration of biomolecular condensates, presenting emerging opportunities to develop therapies targeting transcriptionally addicted malignancies. In this study, we discovered that cyclin-dependent kinase 12 (CDK12), a key regulator of transcription elongation, formed dynamic liquid-like droplets at nuclear speckles, a process enhanced by high CDK12 expression and CDK12 inhibitor exposure. Notably, CDK12 condensation limited sensitivity to kinase inhibition, potentially through enhanced cyclin K recruitment and local enrichment of CDK12-cyclin K complexes. To disrupt these assemblies, we rationally designed HQY1428, a potent modular cyclin K molecular glue degrader that coupled CDK12-binding and DDB1-engaging moieties. Structural and biochemical analyses showed that HQY1428 promoted CDK12-DDB1 association and rapid cyclin K degradation, accompanied by CDK12 depletion and loss of CDK12 condensates. Functionally, HQY1428 treatment perturbed oncogenic transcription and suppressed ovarian cancer growth in vitro and in vivo. Given the frequent genomic co-amplification of *CDK12* and *ERBB2* across diverse human cancers, we showed that combining HQY1428 with HER2 inhibitors provided synergistic therapeutic benefits. Collectively, our findings establish CDK12 condensation at nuclear speckles as an exploitable vulnerability and support targeting CDK12 condensates with cyclin K molecular glue degraders.

## Introduction

Cumulative evidence implicates dysregulated transcription cycles in human cancer and underscores the enormous opportunities of therapeutically targeting the transcriptional machinery (1–3). Many of its components, including RNA polymerase II (RNAPII), transcription factors, mediators, and coactivators, form biomolecular condensates within the nucleus to orchestrate gene expression (4–7). Previous research has primarily focused on the dynamic behaviors and regulatory mechanisms underlying these membraneless compartments assembled through liquid-liquid phase separation (LLPS) (8,9). For example, the intrinsically disordered sequence at the carboxy-terminal domain (CTD) of RNAPII facilitates its clustering and phase separation (10,11). CTD phosphorylation disrupts the RNAPII liquid droplets, allowing multivalent interactions with other elements to drive the transition from transcription initiation to co-transcriptional splicing (12). Recent studies have begun to reveal the functional role of LLPS in partitioning cancer therapeutics, as exemplified by selective accumulation of antineoplastic compounds in specific transcriptional condensates that influences target engagement (13–15). These observations highlight the need to further understand how the condensate organization of transcriptional regulators affects their responses to pharmacological perturbation (16,17).

CDK12 is a pivotal cyclin-dependent kinase that regulates transcriptional cycles in both physiological and pathological contexts (18). CDK12 partners with cyclin K and phosphorylates the serine 2 (S2) residues of tandem heptapeptide repeats (YSPTSPS) in the RNAPII CTD to promote transcription elongation (19,20). Genetically, *CDK12* resides on a chromosome 17q locus proximal to *ERBB2* (encoding HER2) and is frequently co-amplified with *ERBB2* across various cancer types (21–26). Therefore, CDK12 is not only an emerging therapeutic target by itself, but also points to a promising avenue for combination therapy in HER2-positive patients (27,28). At the subcellular level, CDK12 is mainly localized to nuclear speckles (29), specialized ribonucleoprotein granules that organize via phase separation and serve as central hubs for nascent RNA processing (30–32). A significant proportion of nuclear speckle constituents contain low-complexity intrinsically disordered regions (IDRs) (33–35). These observations raise the important question of whether CDK12 forms liquid-like condensates to exert its function at nuclear speckles.

CDK12 as a serine/threonine kinase has been successfully blocked by covalent inhibitors developed by our group and others. However, pharmacokinetic limitations and tolerability concerns often complicate the clinical translation of these compounds (36–40). An alternative and possibly more advantageous strategy is provided by the recent introduction of targeted protein degradation mediated by proteolysis-targeting chimeras (PROTACs) or molecular glue degraders (MGDs) (41–43). Compared with heterobifunctional PROTACs, which consist of two separate ligands and a conjugating linker, monovalent MGDs with generally smaller and more rigid scaffolds have garnered considerable interest owing to their drug-like properties (44). Notably, a spectrum of unexpectedly diverse cyclin K MGDs, e.g., CR8 and HQ461, are serendipitously discovered and act as chemical inducers of proximity between the CDK12-cyclin K complex and the DDB1-CUL4-RBX1 E3 ubiquitin ligase (45–48). Here, we adopted a modular approach and rationally designed a new class of molecular glue degraders by incorporating CDK12-binding and DDB1-engaging moieties. The resulting lead molecule, HQY1428, promoted protein interaction between CDK12 and DDB1, accelerated CDK12-cyclin K degradation, and disrupted CDK12 condensation at nuclear speckles, thereby conferring potent single-agent antitumor activity and synergistic efficacy with HER2 inhibitors. These findings establish a strategy to target CDK12 condensates through cyclin K degradation and support its application in *ERBB2*-amplified cancers.

## Results

### CDK12 forms dynamic condensates at nuclear speckles

Bioinformatics analysis using the Predictor of Natural Disordered Regions (PONDR) and IUPred3 (Figure 1A), together with AlphaFold2 structural predictions (Figure 1B), indicated that both CDK12 and cyclin K contain extensive intrinsically disordered regions (IDRs), suggesting their potential to undergo phase separation. To test this possibility experimentally, we stably expressed monomeric enhanced green fluorescent protein (mEGFP)-tagged CDK12 (CDK12-mEGFP) and cyclin K (cyclin K-mEGFP) in SKOV3 cells. Live-cell fluorescence microscopy revealed that only CDK12-mEGFP formed discrete nuclear puncta, whereas cyclin K-mEGFP exhibited diffuse nuclear distribution (Figure 1C). Treatment with 1,6-hexanediol (1,6-hex), a chemical disruptor of hydrophobic interactions, significantly reduced the number of CDK12-mEGFP puncta (Figure 1D). Fluorescence recovery after photobleaching (FRAP) experiments demonstrated that CDK12-mEGFP puncta rapidly recovered fluorescence within seconds (Figure 1E; Supplementary Figure 1A), supporting their dynamic liquid-like properties. In line with previous reports (29), CDK12-mEGFP puncta prominently colocalized with the nuclear speckle marker SC35 (Figure 1F). To assess endogenous CDK12 distribution, we analyzed its protein levels in a panel of ovarian cancer cell lines (Supplementary Figure 1B). Immunofluorescence staining revealed that endogenous CDK12 also formed distinct nuclear puncta in CDK12-high cell lines (SKOV3, COV362, HEY, OVCA420), while puncta were less apparent in CDK12-low cell lines (TOV-21G, OVCAR3, DOV13, OVCA429) (Supplementary Figure 1C), suggesting an association between CDK12 abundance and puncta accumulation.

**Figure 1.**
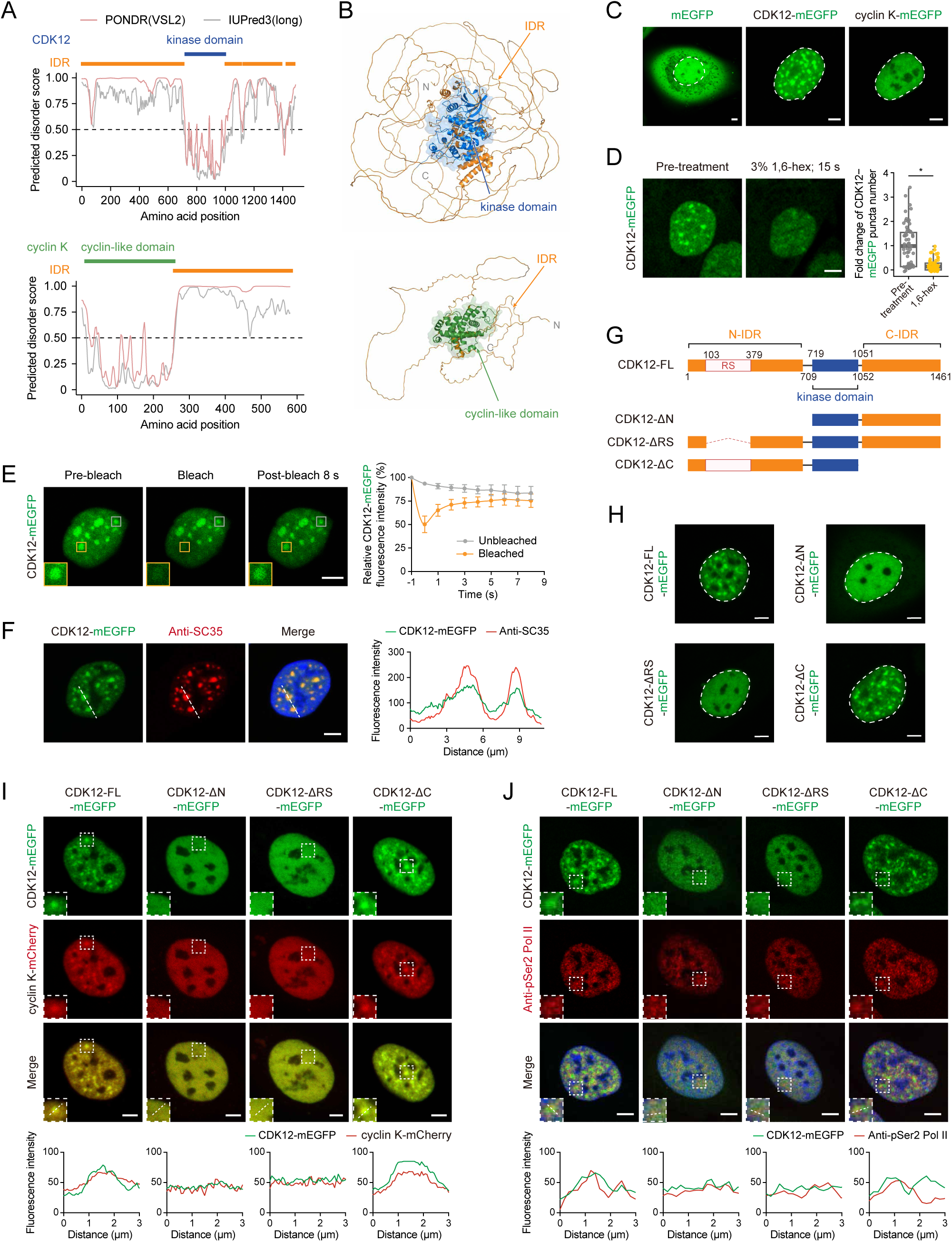
CDK12 forms dynamic condensates at nuclear speckles. A. Graphical depiction of the intrinsically disordered regions (IDRs) of CDK12 and cyclin K, as predicted using the Predictor of Natural Disordered Regions (PONDR) VSL2 and IUPred3 (long mode). Orange bars designate IDRs. The kinase domain of CDK12 and the cyclin-like domain of cyclin K are highlighted with blue and green bars, respectively. B. The protein structures of CDK12 and cyclin K predicted by AlphaFold2, with IDRs and structured domains highlighted. C. Representative live-cell fluorescence images of SKOV3 cells overexpressing mEGFP, CDK12-mEGFP, or cyclin K-mEGFP. Cell nuclei are outlined with dashed white lines. Scale bars, 5 μm. D. Representative images of SKOV3 cells overexpressing CDK12-mEGFP before and after treatment with 3% 1,6-hexanediol (1,6-hex) for 15 s. The box plot shows the fold change of the number of CDK12-mEGFP puncta. Each point represents a single cell. \**P* < 0.05, two-sided Student’s *t*-test. Scale bar, 5 μm. E. Representative images of the fluorescence recovery after photobleaching (FRAP) experiment in SKOV3 cells overexpressing CDK12-mEGFP. The yellow and gray squares indicate the bleached puncta and the unbleached control puncta, respectively. The yellow-boxed region is enlarged in the inset. The relative fluorescence intensities of the bleached area and the unbleached control are quantified and plotted. Data are presented as mean ± s.d. (n = 3). Scale bar, 5 μm. F. Representative immunofluorescence images of CDK12-mEGFP (green) and SC35 (red) in SKOV3 cells. Cell nuclei were counterstained with DAPI (blue). The fluorescence intensity profile along the indicated line is shown on the right. Scale bar, 5 μm. G. Schematic representation of full-length (FL) CDK12 and the indicated deletion mutants. N-IDR, the N-terminal IDR. C-IDR, the C-terminal IDR. RS, the arginine/serine-rich domain. H. Representative images of the indicated mEGFP-tagged CDK12 constructs (green) overexpressed in SKOV3 cells with endogenous *CDK12* knockout. Cell nuclei are outlined with dashed white lines. Scale bars, 5 μm. I. Representative images of the indicated mEGFP-tagged CDK12 constructs (green) co-expressed with cyclin K-mCherry (red) in SKOV3 cells with endogenous *CDK12* knockout. Boxed regions are enlarged in the insets. Fluorescence intensity profiles are plotted below. Scale bars, 5 μm. J. Representative immunofluorescence images of the indicated mEGFP-tagged CDK12 constructs (green) and Ser2-phosphorylated RNA polymerase II (pSer2-Pol II) (red) in SKOV3 cells with endogenous *CDK12* knockout. Cell nuclei were counterstained with DAPI (blue). Boxed regions are enlarged in the insets. Fluorescence intensity profiles are plotted below. Scale bars, 5 μm.

CDK12 contains two major predicted IDRs with high PONDR scores (>0.5). To define the regions required for CDK12 condensate formation, we compared full-length CDK12 (CDK12-FL) with mutants lacking the N-terminal IDR (CDK12-ΔN), the arginine/serine-rich (RS) domain (CDK12-ΔRS), or the C-terminal IDR (CDK12-ΔC) (Figure 1G; Supplementary Figure 1D). Deletion of either the N-terminal IDR or the RS-rich domain markedly impaired the formation of CDK12 puncta, whereas deletion of the C-terminal IDR retained a punctate distribution (Figure 1H).

Interestingly, co-expression of CDK12-mEGFP increased cyclin K puncta intensity (Supplementary Figure 1E). Moreover, cyclin K was enriched in CDK12 puncta at nuclear speckles in cells expressing CDK12-FL or CDK12-ΔC, but remained largely diffuse in those expressing CDK12-ΔN or CDK12-ΔRS (Figure 1I). Ser2-phosphorylated RNA polymerase II (pSer2-Pol II) localized near the periphery of CDK12-FL and CDK12-ΔC condensates, whereas this spatial organization was not apparent with CDK12-ΔN or CDK12-ΔRS (Figure 1J). Given the close spatial association between pSer2-Pol II clusters and nascent RNA (49,50), these observations suggest that CDK12 condensates are positioned near sites of active transcription. Together, these results support a model in which CDK12 condensation promotes cyclin K partitioning and facilitates the assembly of transcriptionally active complexes at nuclear speckles.

### CDK12 condensation limits sensitivity to kinase inhibition

Given the emerging data highlighting the interplay between phase-separated condensates and small-molecule compounds (13–15), we sought to determine how CDK12-targeting compounds affect CDK12 condensates and whether CDK12 condensation influences drug response. Treatment with the covalent CDK12/13 kinase inhibitor THZ531 (37) increased CDK12 puncta accumulation in cells expressing CDK12-FL or CDK12-ΔC, whereas CDK12-ΔN and CDK12-ΔRS remained largely diffuse (Figure 2A). Notably, cells expressing CDK12-ΔN or CDK12-ΔRS were more sensitive to THZ531 than those expressing CDK12-FL or CDK12-ΔC (Figure 2B). The enrichment of cyclin K in CDK12 condensates (Figure 1I) could help explain these differences in THZ531 sensitivity. Indeed, THZ531-induced accumulation of CDK12-FL puncta was accompanied by enhanced recruitment of cyclin K into these condensates (Figure 2C), consistent with increased local concentration of CDK12-cyclin K complexes that might counteract the effects of THZ531 treatment. By contrast, cyclin K remained largely diffuse in cells expressing CDK12-ΔRS (Figure 2C). To assess whether the effects on CDK12 condensation extended beyond THZ531, we treated SKOV3 cells with additional CDK12-binding kinase inhibitors and degraders in the presence of MLN4924, a NEDD8-activating enzyme inhibitor which blocks cullin-RING ligase activity (51). Under these conditions, similar increases in CDK12 puncta accumulation at SC35-marked nuclear speckles were observed following treatment with the tested compounds (Figure 2D). Together, these findings suggest that pharmacological targeting of CDK12 induces its condensation and promotes cyclin K enrichment at nuclear speckles, potentially limiting inhibitor efficacy.

**Figure 2.**
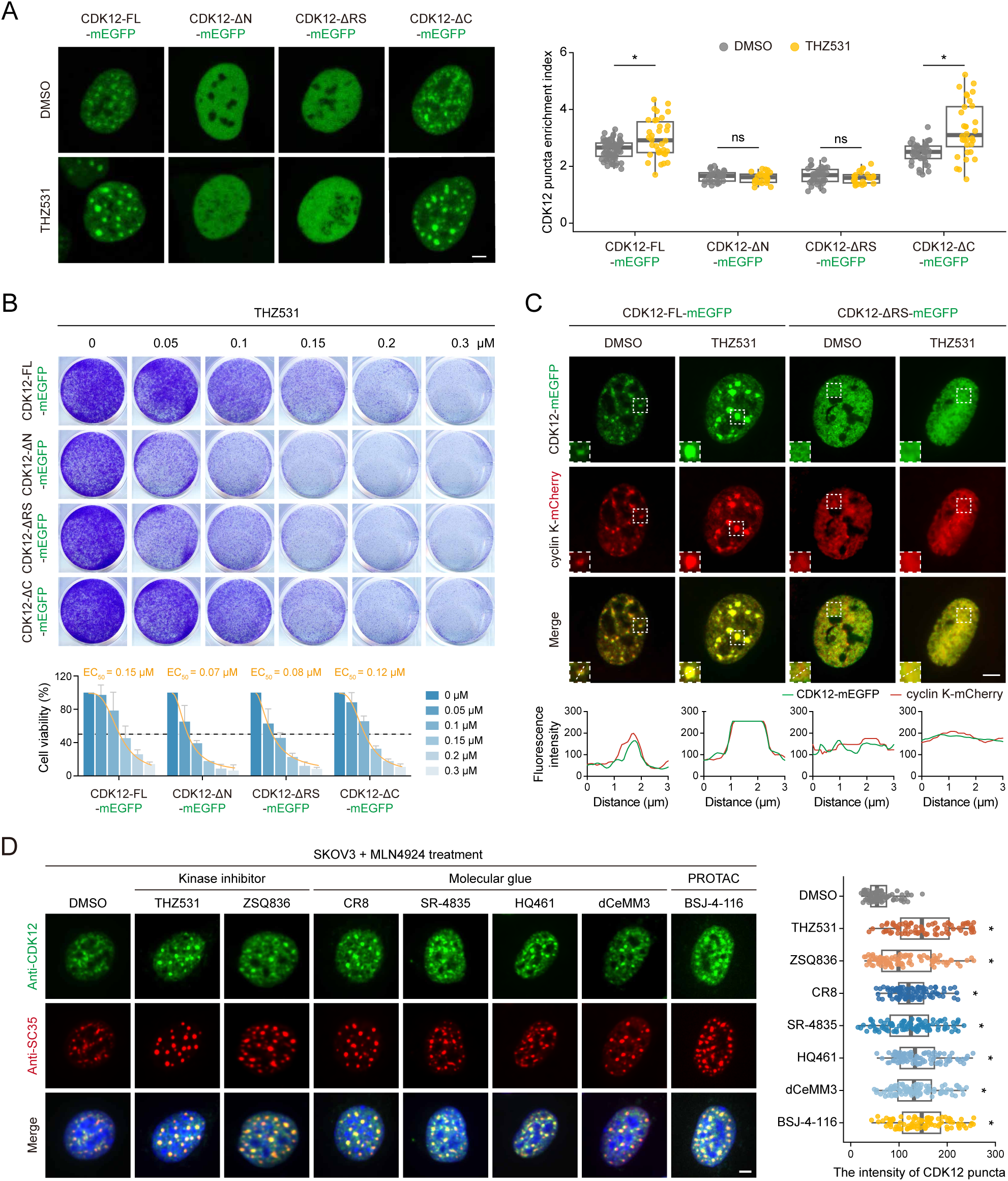
CDK12 condensation limits sensitivity to kinase inhibition. A. Representative fluorescence images of the indicated mEGFP-tagged CDK12 constructs (green) overexpressed in SKOV3 cells with endogenous *CDK12* knockout after treatment with DMSO or THZ531 (1 µM) for 24 h. Quantification of the CDK12 puncta enrichment index is plotted on the right. Box plot shows CDK12 puncta enrichment index for each group. \**P* < 0.05; ns, not significant, Kruskal-Wallis test with Holm correction. Scale bar, 5 μm. B. SKOV3 cells with endogenous *CDK12* knockout expressing mEGFP-tagged CDK12 constructs were treated with various concentrations of THZ531 for 5 days. Cell viability was estimated by densitometric analysis of crystal violet staining. Data are presented as mean ± s.d. (n = 3). The half-maximal effective concentration (EC50) values are shown. C. Representative fluorescence images of CDK12-FL-mEGFP or CDK12-ΔRS-mEGFP (green) with cyclin K-mCherry (red) in SKOV3 cells with endogenous *CDK12* knockout after treatment with DMSO or THZ531 (1 µM) for 24 h. Boxed regions are enlarged in the insets. Fluorescence intensity profiles along the indicated lines are shown below. Scale bar, 5 μm. D. Representative immunofluorescence images of endogenous CDK12 (green) and SC35 (red) in SKOV3 cells. Cells were pretreated with MLN4924 (0.5 μM) for 2 h, and then treated for an additional 24 h with DMSO, THZ531 (1 μM), ZSQ836 (1 μM), CR8 (2 μM), SR-4835 (2 μM), HQ461 (10 μM), dCeMM3 (20 μM), or BSJ-4-116 (1 μM) in the continued presence of MLN4924 (0.5 μM). Cell nuclei were counterstained with DAPI (blue). Box plot shows the intensity of CDK12 puncta for each group. \**P* < 0.05, Kruskal-Wallis test with Holm correction. Scale bar, 5 μm.

### Rational design and characterization of potent cyclin K molecular glue degraders

We reasoned that cyclin K degradation induced by molecular glues such as CR8 could dismantle the CDK12-cyclin K assemblies that persisted during kinase inhibition and that more efficient protein depletion would improve drug efficacy (Supplementary Figure 2A). We therefore adopted a modular design strategy to develop next-generation cyclin K molecular glue degraders. The CDK12-binding module combined a head moiety derived from TL12-186 (52), a pan-kinase PROTAC known to degrade CDK12, with a piperidine-containing linker inspired by THZ531 (37), a covalent CDK12/13 inhibitor whose linker configuration contributed to its selectivity over CDK7/9. A terminal phenyl group was selected based on evaluation of the DDB1-engaging moieties of CR8 and related analogs, including UB-007 (53). Assembly of these subunits yielded ZSQ253 (Figure 3A; Supplementary Figure 2B). ZSQ253 effectively depleted cyclin K in SKOV3 cells, and this effect was blocked by the proteasome inhibitor PS341 but not by chloroquine, supporting proteasome-dependent degradation (Supplementary Figure 2C). To further improve its pharmacological properties and degradation potency, we pursued optimization of its tail moiety. Molecular docking was performed based on the CR8-bound CDK12-cyclin K-DDB1 co-crystal structure (PDB: 6TD3), which predicted a plausible binding mode for ZSQ253 with a docking score of-13.90 (Supplementary Figure 2D). In this model, ZSQ253 occupied the CDK12 ATP-binding pocket similarly to CR8, with its aminopyrimidine moiety forming hydrogen bonds with the hinge region and its phenyl tail engaging DDB1 residues via cation-π interactions. Given the unoccupied space surrounding the tail moiety, a series of analogs with tail modifications were designed and evaluated by docking to exploit potential additional interactions. Notably, HQY1428, featuring a 3-amino-5-fluorophenyl tail, exhibited a more favorable docking score (-14.94) and was predicted to form additional contacts with Ile733 of CDK12 and Phe949 of DDB1, potentially further stabilizing the ternary complex. In contrast, replacing the phenyl tail of ZSQ253 with a methyl group removed the aromatic DDB1-engaging moiety and yielded a less favorable docking score (-7.17). The resulting compound, ZSQ2549, served as a non-degrading control that retained CDK12 kinase-inhibitory activity (Figure 3A; Supplementary Figure 2D).

**Figure 3.**
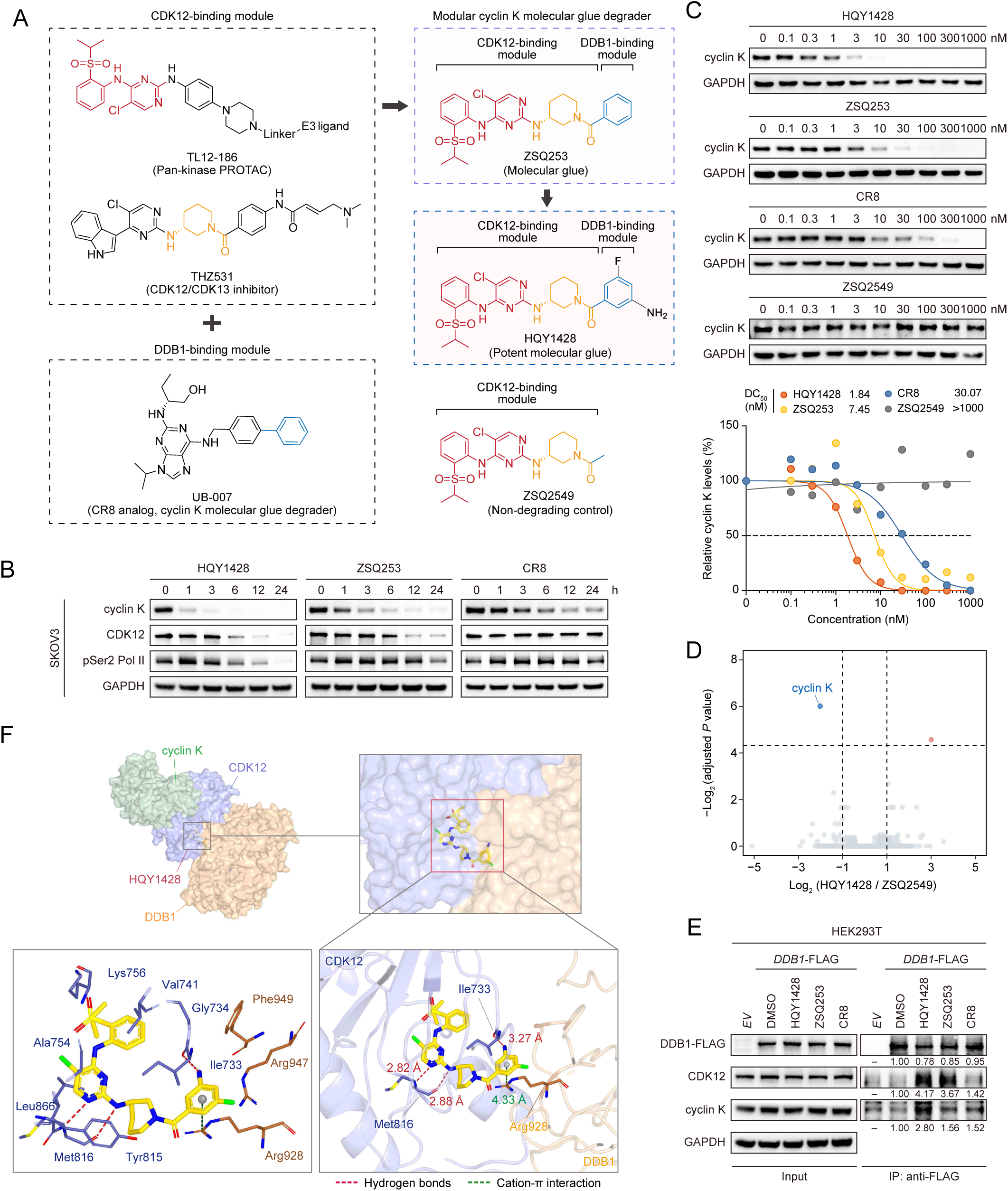
Rational design and characterization of potent cyclin K molecular glue degraders. A. Schematic illustration of the rationale for the modular design of the cyclin K molecular glue degraders ZSQ253 and HQY1428. The CDK12-binding module was derived from TL12-186 (pan-kinase PROTAC) and THZ531 (CDK12/13 inhibitor). The DDB1-binding module was derived from UB-007, a CR8 analog. The chemical structure of the non-degrading control compound ZSQ2549 is also shown. B. Immunoblotting analysis of cyclin K, CDK12, and Ser2-phosphorylated RNA polymerase II (pSer2-Pol II) in SKOV3 cells treated with HQY1428 (0.1 μM), ZSQ253 (0.1 μM), or CR8 (0.1 μM) over a time course. GAPDH was used as the loading control. C. Immunoblotting analysis of cyclin K in SKOV3 cells treated with the indicated concentrations of HQY1428, ZSQ253, CR8 or ZSQ2549. GAPDH was used as the loading control. Quantification of cyclin K protein levels is shown below. Curves were fitted to estimate the half-maximal degradation concentration (DC50). D. Whole-proteome quantification of SKOV3 cells treated with HQY1428 (0.1 μM) or ZSQ2549 (0.1 μM) for 4 h (n = 3 per group). Dashed lines indicate adjusted *P* = 0.05 and log₂ fold change = ±1. E. Co-immunoprecipitation analysis of the association of CDK12 and cyclin K with DDB1 in HEK293T cells transfected with empty vector (*EV*) or FLAG-tagged *DDB1*. Cells were pretreated with MLN4924 (0.5 μM) for 2 h, followed by treatment with DMSO, HQY1428 (0.1 μM), ZSQ253 (0.1 μM), or CR8 (0.1 μM) for 1 h. Input lysates and anti-FLAG immunoprecipitates were analyzed by immunoblotting with the indicated antibodies. GAPDH and DDB1-FLAG served as loading controls for the input and immunoprecipitated fractions, respectively. F. Cryo-electron microscopy (cryo-EM) structure of the HQY1428-bound CDK12-cyclin K-DDB1 complex, determined at an overall resolution of 3.76 Å. CDK12, cyclin K and DDB1 are shown in purple, green and orange, respectively. HQY1428 is displayed as yellow sticks and occupies the interface between CDK12 and DDB1. Enlarged views highlight the HQY1428-binding pocket and its interactions with surrounding residues. Red and green dashed lines indicate hydrogen bonds and a cation-π interaction, respectively.

Guided by molecular docking, HQY1428 and ZSQ2549 were synthesized and evaluated. Compared to ZSQ253, HQY1428 exhibited improved aqueous solubility, attributable to its polar and salifiable amino group, and greater metabolic stability in mouse liver microsomes (Supplementary Figure 2E). Western blot analysis demonstrated that HQY1428 induced more rapid cyclin K degradation than ZSQ253 or CR8 in SKOV3 cells, accompanied by CDK12 depletion and reduced RNA polymerase II CTD Ser2 phosphorylation (Figure 3B). At 100 nM treatment, the apparent cyclin K depletion half-times were 0.40, 1.36, and 2.69 h for HQY1428, ZSQ253, and CR8, respectively (Supplementary Figure 2F). The superior degradation potency of HQY1428 was further corroborated by dose-response titrations, yielding a half-maximal degradation concentration (DC50) value of 1.84 nM, approximately 4-fold and 16-fold lower than those of ZSQ253 and CR8, respectively (Figure 3C). In contrast, ZSQ2549 failed to induce cyclin K degradation even at 1 μM. Furthermore, global proteomic profiling via mass spectrometry of SKOV3 cells treated with HQY1428 or ZSQ2549 for 4 h revealed cyclin K as the most significantly downregulated protein, establishing HQY1428 as a highly potent and selective cyclin K degrader (Figure 3D; Supplementary Table 1).

We next validated the molecular glue degrader mechanism for HQY1428. As expected, treatment with MLN4924 completely blocked cyclin K degradation induced by HQY1428, ZSQ253, or CR8, confirming a cullin-RING ligase-dependent mechanism (Supplementary Figure 2G). To further delineate the required E3 ligase machinery, we generated individual knockouts of core components of the DDB1-CUL4-RBX1 complex in SKOV3 cells. Deletion of *DDB1* or *RBX1* abrogated cyclin K degradation elicited by all three compounds (Supplementary Figure 2H). In contrast, individual knockout of *CUL4A* or *CUL4B* did not prevent cyclin K degradation, likely due to functional redundancy between the two paralogs (54). Consistent with these data, co-immunoprecipitation assays revealed that HQY1428 enhanced the association of CDK12-cyclin K with DDB1 to a greater extent than ZSQ253 or CR8 (Figure 3E).

To elucidate the structural basis for the superior ability of HQY1428 to promote ternary complex formation, we sought to determine the cryo-electron microscopy (cryo-EM) structure of the HQY1428-bound CDK12-cyclin K-DDB1 complex (Supplementary Figure 3A). We obtained a reconstruction at an overall resolution of 3.76 Å (Supplementary Figure 3B). The structural model illustrated the positioning of HQY1428 at the CDK12-DDB1 interface (Supplementary Figure 3C), with hydrogen-bonding contacts involving the backbone of Met816 in the CDK12 hinge region and a cation-π interaction between its terminal phenyl ring and Arg928 on DDB1, consistent with our molecular docking predictions (Figure 3F). Notably, the amino group of HQY1428 was positioned approximately 3.27 Å from the backbone carbonyl oxygen of Ile733 on CDK12, suggestive of an additional hydrogen bond. Furthermore, the fluorine atom of HQY1428 engaged surrounding DDB1 residues, such as Arg928 and Arg947, through hydrophobic interactions. These structural features provide a rationale for HQY1428-mediated DDB1 recruitment and, together with the biochemical and genetic evidence, support a DDB1-CUL4-RBX1-dependent mechanism of cyclin K degradation.

### HQY1428 disrupts CDK12 condensates and inhibits tumor cell viability

Given the significantly enhanced robustness of cyclin K degradation by HQY1428, we next investigated whether this translated into more efficient disruption of CDK12 condensates. Immunofluorescence staining showed a more rapid loss of CDK12 puncta at nuclear speckles following HQY1428 treatment (Figure 4A; Supplementary Figure 4A), which correlated with stronger growth inhibition compared to ZSQ253 or CR8 (Supplementary Figure 4B). The growth-inhibitory effects of HQY1428, ZSQ253, and CR8 were all mitigated by co-treatment with MLN4924 (Figure 4B) or by deletion of *DDB1* (Figure 4C). Specifically, MLN4924 increased the EC₅₀ of HQY1428 from 0.001 to 0.20 μM, an approximately 200-fold shift, compared with 72.5-fold and 16.9-fold shifts for ZSQ253 and CR8, respectively. *DDB1* deletion also markedly reduced sensitivity to HQY1428. In contrast, these perturbations had minimal impact on the response to the non-degrading kinase inhibitor ZSQ2549. When further benchmarked against other kinase inhibitors and degraders, HQY1428 exhibited the greatest growth-inhibitory potency in SKOV3 cells (Figure 4D) and induced the most pronounced cyclin K depletion (Figure 4E). Collectively, these results establish HQY1428 as a highly efficient and specific cyclin K molecular glue degrader that facilitates DDB1 recruitment, accelerates cyclin K depletion, disrupts CDK12 condensates, and exhibits robust antiproliferative activity. The rapid degradation and associated loss of CDK12 condensates likely contribute to the enhanced cellular drug sensitivity.

**Figure 4.**
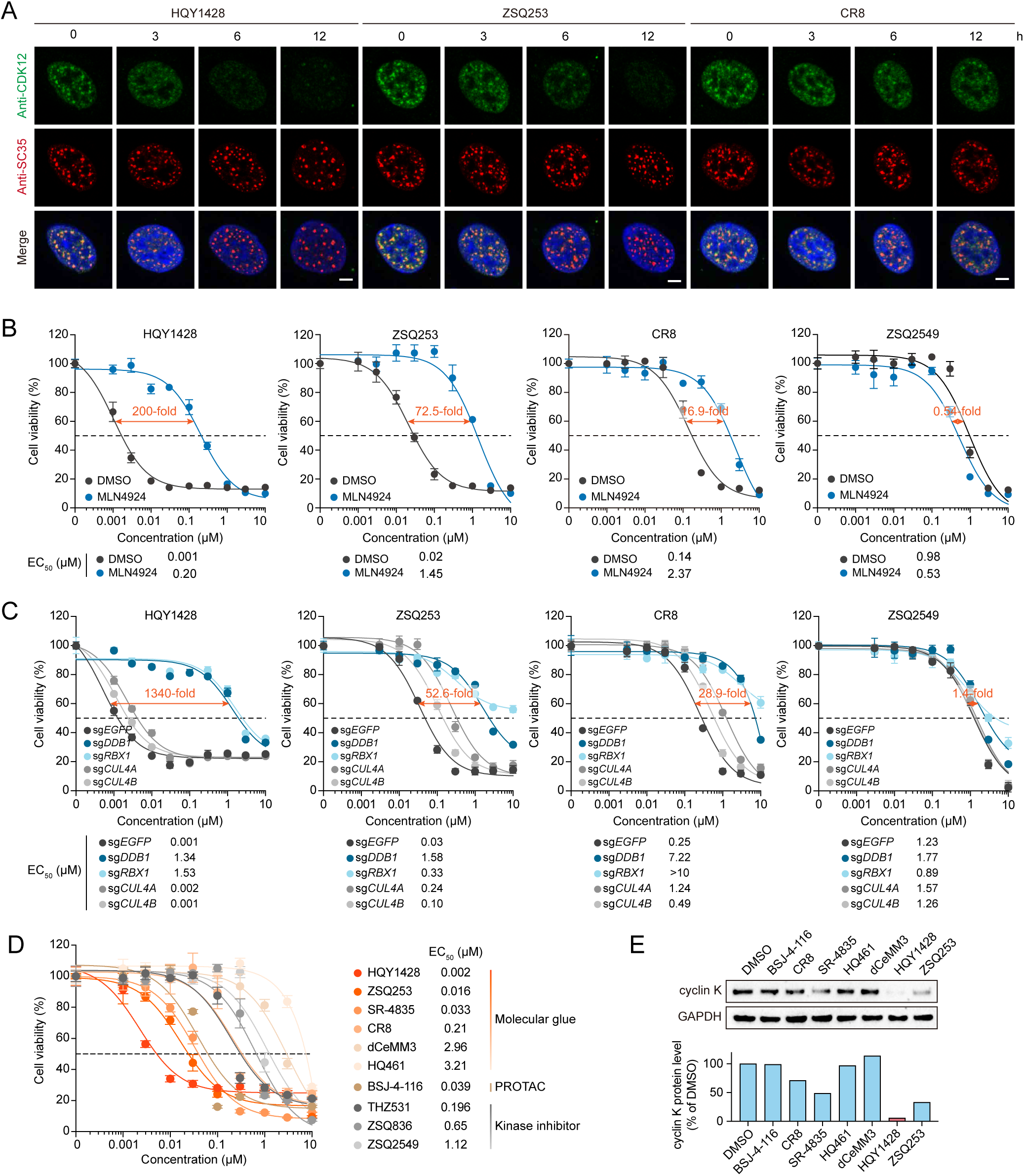
HQY1428 disrupts CDK12 condensates and inhibits tumor cell viability. A. Representative immunofluorescence images of CDK12 (green) and SC35 (red) in SKOV3 cells treated with HQY1428 (0.1 μM), ZSQ253 (0.1 μM), or CR8 (0.1 μM) over a time course. Cell nuclei were counterstained with DAPI (blue). Scale bars, 5 μm. B. SKOV3 cells were pretreated with DMSO or MLN4924 (0.5 μM) for 2 h and then treated with the indicated compounds at various concentrations in the continued presence or absence of MLN4924 for 72 h. Cell viability was measured using the Cell Counting Kit-8 (CCK-8) assay. Data are presented as mean ± s.d. (n = 3). EC50 values and fold changes in EC₅₀ upon MLN4924 treatment relative to the DMSO control are shown. C. SKOV3 cells with *DDB1*, *RBX1*, *CUL4A*, or *CUL4B* knockout were treated with various concentrations of HQY1428, ZSQ253, CR8, or ZSQ2549 for 72 h. Cell viability was measured using the CCK-8 assay. Data are presented as mean ± s.d. (n = 3). EC50 values and fold changes in EC₅₀ for *DDB1*-depleted cells relative to sg*EGFP* control cells are shown. D. SKOV3 cells were treated with the indicated compounds at various concentrations for 72 h. Cell viability was measured using the CCK-8 assay. Data are presented as mean ± s.d. (n = 3). EC50 values are shown. E. Immunoblotting analysis of cyclin K in SKOV3 cells treated with the indicated compounds (0.1 µM each) for 1 h. GAPDH was used as the loading control. Quantification of cyclin K protein levels relative to DMSO is shown below.

### Transcriptional responses to cyclin K molecular glue degraders

To characterize the transcriptomic effects of cyclin K molecular glue degraders, we performed RNA sequencing (RNA-seq) on drug-exposed SKOV3 cells (Supplementary Figure 5A). Following brief treatment with CR8 (Supplementary Table 2), ZSQ253 (Supplementary Table 3), or HQY1428 (Supplementary Table 4), we observed widespread alterations in gene expression, with HQY1428 exhibiting the most pronounced effects (Figure 5A). Differentially expressed genes (DEGs) were predominantly downregulated across all three treatments, with HQY1428 producing the largest number of DEGs (Figure 5B). A significant positive correlation of transcriptome-wide changes was found across the three compounds (Supplementary Figure 5B), indicating broadly concordant responses. Gene set enrichment analysis (GSEA) pinpointed suppressed pathways related to DNA repair, DNA replication, cell cycle, RNA metabolism, and transcriptional regulation (Figure 5C), in keeping with the antitumor activity. Likewise, Cytoscape analysis revealed multiple disrupted oncogenic pathways, especially those involved in DNA damage response (DDR) (Supplementary Figure 5C). Accordingly, the expression of 276 DDR genes curated by TCGA (55) was disproportionately suppressed following all three treatments, with the most negative normalized enrichment score observed for HQY1428 (Figure 5D). Consistent with this finding, multiple core DDR genes involved in distinct DNA repair pathways were downregulated (Supplementary Figure 5D).

**Figure 5.**
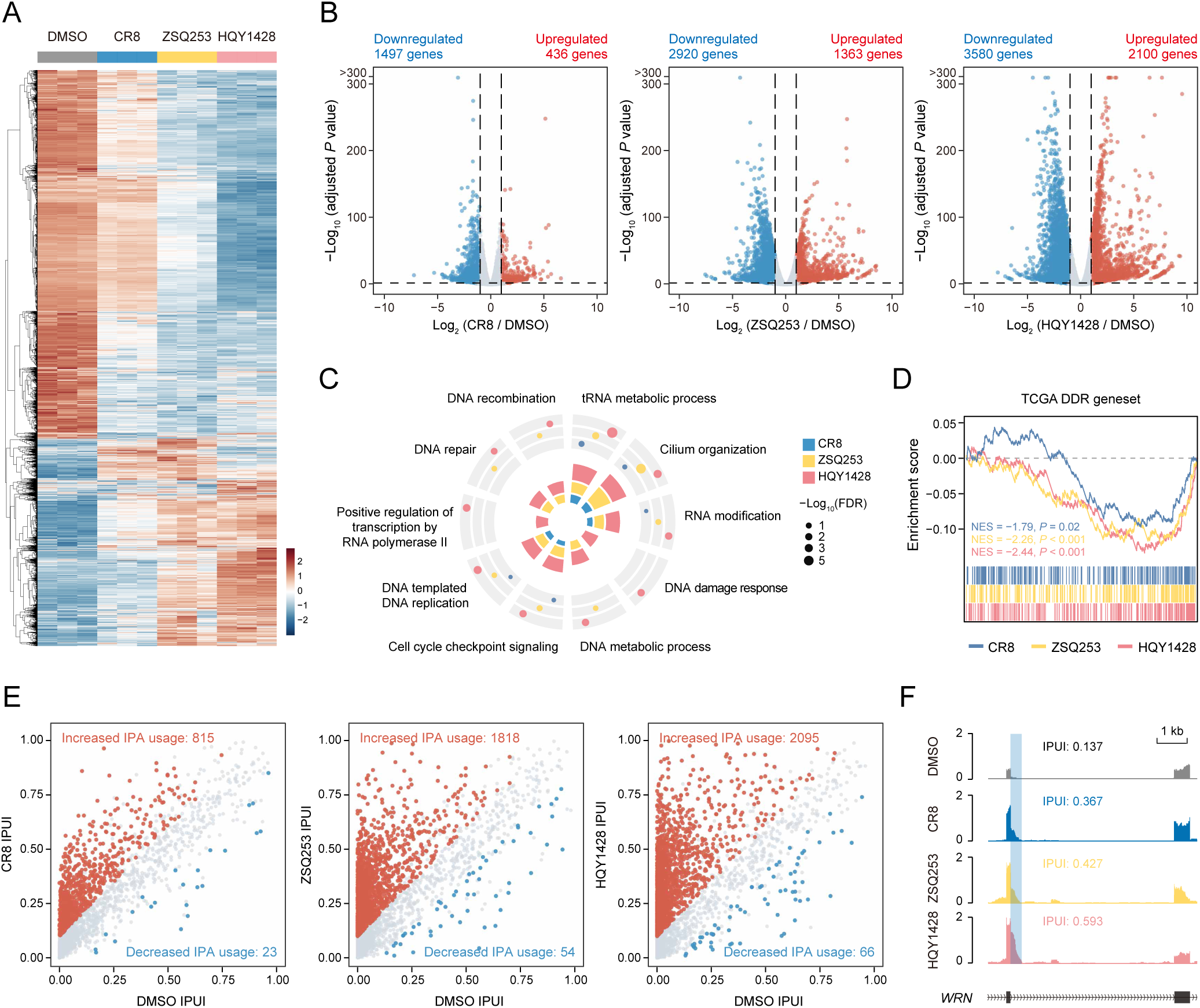
Transcriptional responses to cyclin K molecular glue degraders. A. Heatmap of differentially expressed genes in SKOV3 cells treated with DMSO, CR8, ZSQ253, or HQY1428. Rows represent genes, and columns represent samples. Colors indicate scaled expression levels. B. Volcano plots showing differentially expressed genes in SKOV3 cells treated with CR8, ZSQ253, or HQY1428 (0.1 µM each) for 6 h relative to DMSO-treated controls. Differentially expressed genes were defined by adjusted P < 0.05 and |log₂ fold change| > 1. C. Circular plot showing gene set enrichment analysis of selected biological processes in SKOV3 cells treated with CR8, ZSQ253, or HQY1428 for 6 h. The radial height of each colored segment in the inner plot indicates the absolute value of the normalized enrichment score. Dot size indicates −log₁₀(FDR). D. GSEA plots showing disproportionate downregulation of a TCGA-defined set of 276 DNA damage repair genes in SKOV3 cells treated with CR8, ZSQ253, or HQY1428 relative to DMSO-treated controls. NES, normalized enrichment score. E. Scatter plots showing differential intronic polyadenylation (IPA) events (adjusted *P* < 0.05 and |IPUI difference| > 0.1) in SKOV3 cells treated with CR8, ZSQ253, or HQY1428 relative to DMSO-treated controls. The numbers of differential IPA events are indicated. F. RNA-seq read density illustrating the relative IPA usage at the *WRN* gene locus in SKOV3 cells treated with DMSO, CR8, ZSQ253, or HQY1428. Intronic poly(A) site usage index (IPUI) values are indicated.

Given that CDK12 suppresses intronic polyadenylation (IPA) to maintain DNA repair gene expression (56), we examined whether cyclin K molecular glue degraders affected IPA usage. CR8, ZSQ253, and HQY1428 increased IPA usage at 815, 1,818, and 2,095 events, respectively, with substantially fewer decreases (Figure 5E; Supplementary Tables 5-7). All three treatments increased IPA usage in 534 shared genes (Supplementary Figure 5E), as exemplified by *WRN*, whose intronic poly(A) site usage index (IPUI) was highest following HQY1428 treatment (Figure 5F). Considering that nuclear speckles serve as central hubs for splicing regulation, we further conducted alternative splicing analysis using the rMATS pipeline (57) and identified 2,045, 3,731, and 5,068 events in response to CR8, ZSQ253, and HQY1428, respectively (Supplementary Figure 5F; Supplementary Tables 8-10). There were 740 differentially spliced genes shared by all three treatments, with HQY1428 affecting the largest number of genes (Supplementary Figure 5G). Taken together, these results indicate that HQY1428 induces dramatic transcriptional alterations encompassing changes in gene expression and RNA processing.

### HQY1428 induces DNA damage and suppresses ovarian cancer growth

Given its enhanced cellular potency, we selected HQY1428 for further evaluation in ovarian cancer models. HQY1428 induced concentration-dependent growth inhibition across a panel of ovarian cancer cell lines (Figure 6A; Supplementary Figure 6A). EdU (5-ethynyl-2’-deoxyuridine) incorporation assays showed that HQY1428 significantly reduced DNA synthesis (Figure 6B). Moreover, HQY1428 treatment induced G2/M arrest (Figure 6C) and cell apoptosis, as evidenced by increased caspase-3/7 activity (Figure 6D; Supplementary Figure 6B) and annexin V staining (Figure 6E). In line with our RNA-seq findings and the established role of CDK12 in maintaining DDR gene expression (56,58), Western blot analysis revealed elevated γH2AX and reduced RAD51 levels, accompanied by cyclin K and CDK12 depletion and reduced RNA polymerase II CTD Ser2 phosphorylation (Figure 6F). Consistently, immunofluorescence staining confirmed a marked increase in nuclear γH2AX foci following HQY1428 treatment, indicating the accumulation of DNA double-strand breaks (Figure 6G). These findings demonstrate that HQY1428 induces DNA damage and exerts pronounced antiproliferative and cytotoxic effects in ovarian cancer cells.

**Figure 6.**
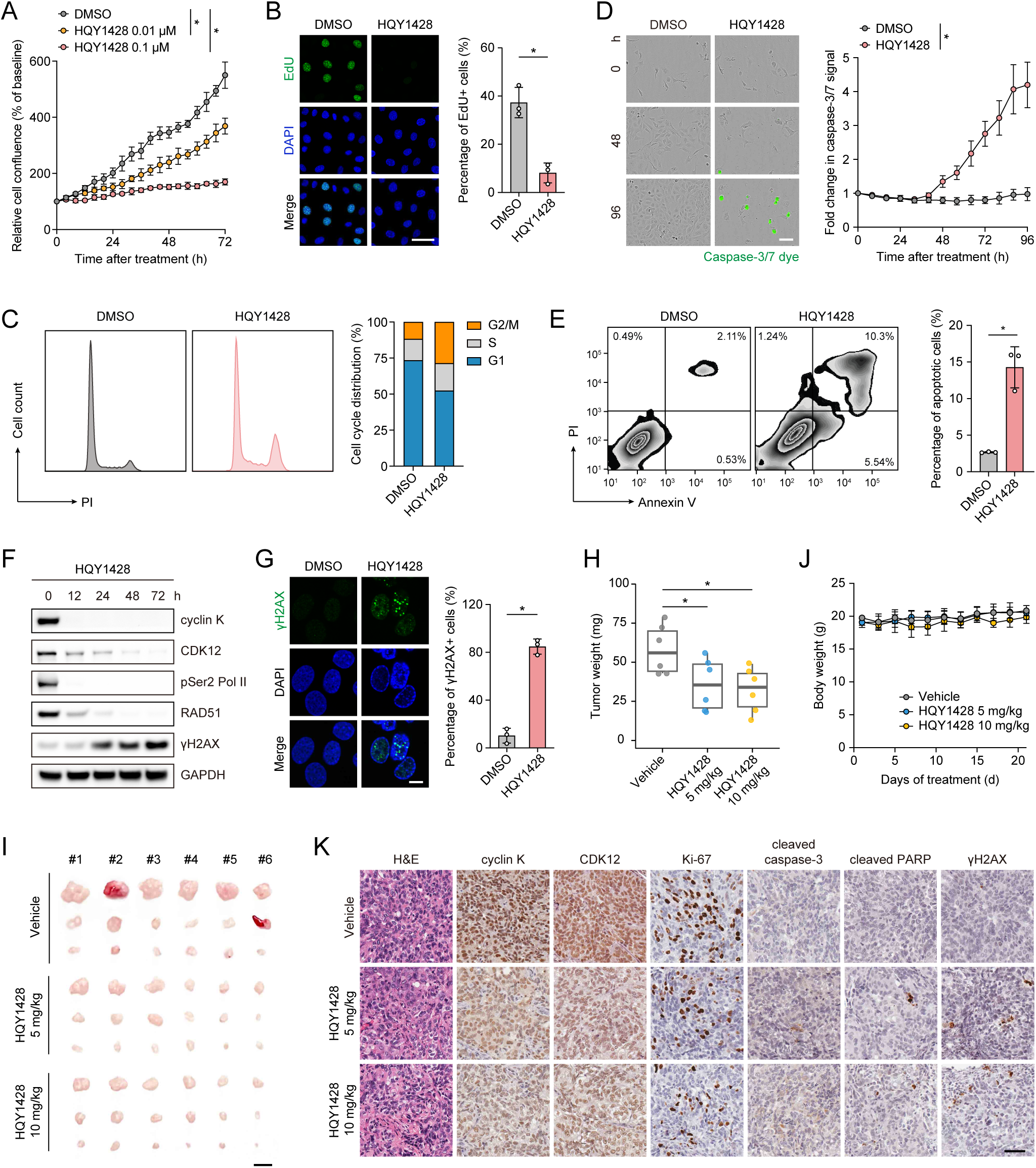
HQY1428 induces DNA damage and suppresses ovarian cancer growth. A. SKOV3 cells were treated with DMSO or the indicated concentrations of HQY1428, and cell growth was monitored for 72 h using the Incucyte live-cell imaging system. Data are presented as mean ± s.d. (n = 4). \**P* < 0.05, two-way ANOVA. B. EdU (green) incorporation assay in SKOV3 cells after treatment with DMSO or HQY1428 (0.1 μM) for 24 h. Cell nuclei were counterstained with DAPI (blue). The percentage of EdU-positive cells is plotted as mean ± s.d. (n = 3). \**P* < 0.05, unpaired Student’s *t*-test. Scale bar, 50 μm. C. Cell cycle analysis by flow cytometry in SKOV3 cells after treatment with DMSO or HQY1428 (0.1 μM) for 48 h. The proportions of cells in G1, S, and G2/M phases are shown on the right. D. Representative images of SKOV3 cells treated with DMSO or HQY1428 (0.1 μM) in the presence of caspase-3/7 green detection reagent. Changes in caspase-3/7 fluorescence over 96 h are shown on the right. Data are presented as mean ± s.d. (n = 4). \**P* < 0.05, two-way ANOVA. Scale bar, 200 μm. E. Flow cytometric analysis of cell death using annexin V/propidium iodide (PI) staining in SKOV3 cells after treatment with DMSO or HQY1428 (0.1 μM) for 72 h. The percentage of apoptotic cells is plotted as mean ± s.d. (n = 3). \**P* < 0.05, unpaired Student’s *t*-test. F. Immunoblotting analysis of the indicated proteins in SKOV3 cells treated with HQY1428 (0.1 μM) over a time course. GAPDH was used as the loading control. G. Representative immunofluorescence images of γH2AX (green) in SKOV3 cells treated with DMSO or HQY1428 (0.1 μM) for 72 h. Cell nuclei were counterstained with DAPI (blue). The percentage of cells containing more than five γH2AX foci is plotted as mean ± s.d. (n = 3). \**P* < 0.05, unpaired Student’s *t*-test. Scale bar, 10 μm. H. Quantification of total intraperitoneal SKOV3 tumor weight following oral administration of vehicle control (2.5% v/v DMSO and 97.5% v/v 30% SBE-β-CD) or HQY1428 (5 or 10 mg/kg/day) for 21 days. The tumor weight for each mouse was calculated by adding the weights of all resectable implants. Each point represents one mouse. \**P* < 0.05, ANOVA followed by Tukey’s post-test. I. Representative images of SKOV3 xenografts from BALB/c nude mice treated with vehicle control or HQY1428 via oral gavage for 21 days. The three largest abdominal masses are shown for each mouse. Scale bar, 5 mm. J. Body weight measurements of BALB/c nude mice during vehicle control or HQY1428 treatment. K. Representative images of hematoxylin and eosin (H&E) and immunohistochemistry (IHC) staining for cyclin K, CDK12, Ki-67, cleaved caspase-3, cleaved PARP, and γH2AX in SKOV3 tumor sections. Scale bar, 50 μm.

To evaluate the in vivo efficacy, HQY1428 was administered by oral gavage at 5 or 10 mg/kg/day in BALB/c nude mice bearing intraperitoneal SKOV3 xenografts. Treatment with HQY1428 resulted in markedly reduced tumor weight at both doses (Figure 6H), with smaller abdominal implants observed at necropsy (Figure 6I). Importantly, no significant loss of body weight (Figure 6J) or overt histological abnormalities in the heart, liver, spleen, lung, and kidney were observed (Supplementary Figure 6C). In agreement with the in vitro data, tumor immunohistochemistry showed decreased levels of cyclin K and CDK12, reduced cell proliferation as indicated by Ki-67 staining, increased cell apoptosis as marked by cleaved caspase-3 and cleaved PARP, and elevated γH2AX-labeled DNA damage (Figure 6K). These results underscore the in vivo antitumor efficacy and favorable safety profile of HQY1428 as a promising cyclin K molecular glue degrader for future development.

### HQY1428 enhances the antitumor activity of HER2 inhibitors

*CDK12* and *ERBB2* are located in close genomic proximity and are frequently co-amplified in many cancer types (21–26). Indeed, the two genes showed concordant copy number patterns across 9,470 TCGA samples (Figure 7A), with *CDK12* amplification occurring almost exclusively in *ERBB2*-amplified tumors (Supplementary Figure 7A). Consequently, their transcript levels were also positively correlated (Supplementary Figure 7B). To corroborate these observations at the protein level, we evaluated a diverse cohort of primary HER2-positive malignancies, including breast, gastric, biliary tract, colorectal, salivary duct, and urothelial cancers (Supplementary Table 11). Immunofluorescence staining revealed prominent nuclear CDK12 staining and distinct puncta at SC35-marked nuclear speckles (Figure 7B; Supplementary Figure 8). Subsequent analysis of the Cancer Cell Line Encyclopedia (CCLE) similarly linked *ERBB2* amplification to *CDK12* amplification (Supplementary Figure 9A) and elevated *CDK12* expression (Supplementary Figure 9B). Accordingly, CDK12 condensates at nuclear speckles were ubiquitously detected in a panel of *ERBB2*-amplified cell lines derived from breast (AU565, SKBR3, BT474), gastric (NCI-N87), esophageal (TE4), lung (Calu-3, NCI-H2170), and ovarian (SKOV3) cancers (Figure 7C; Supplementary Figure 9C).

**Figure 7.**
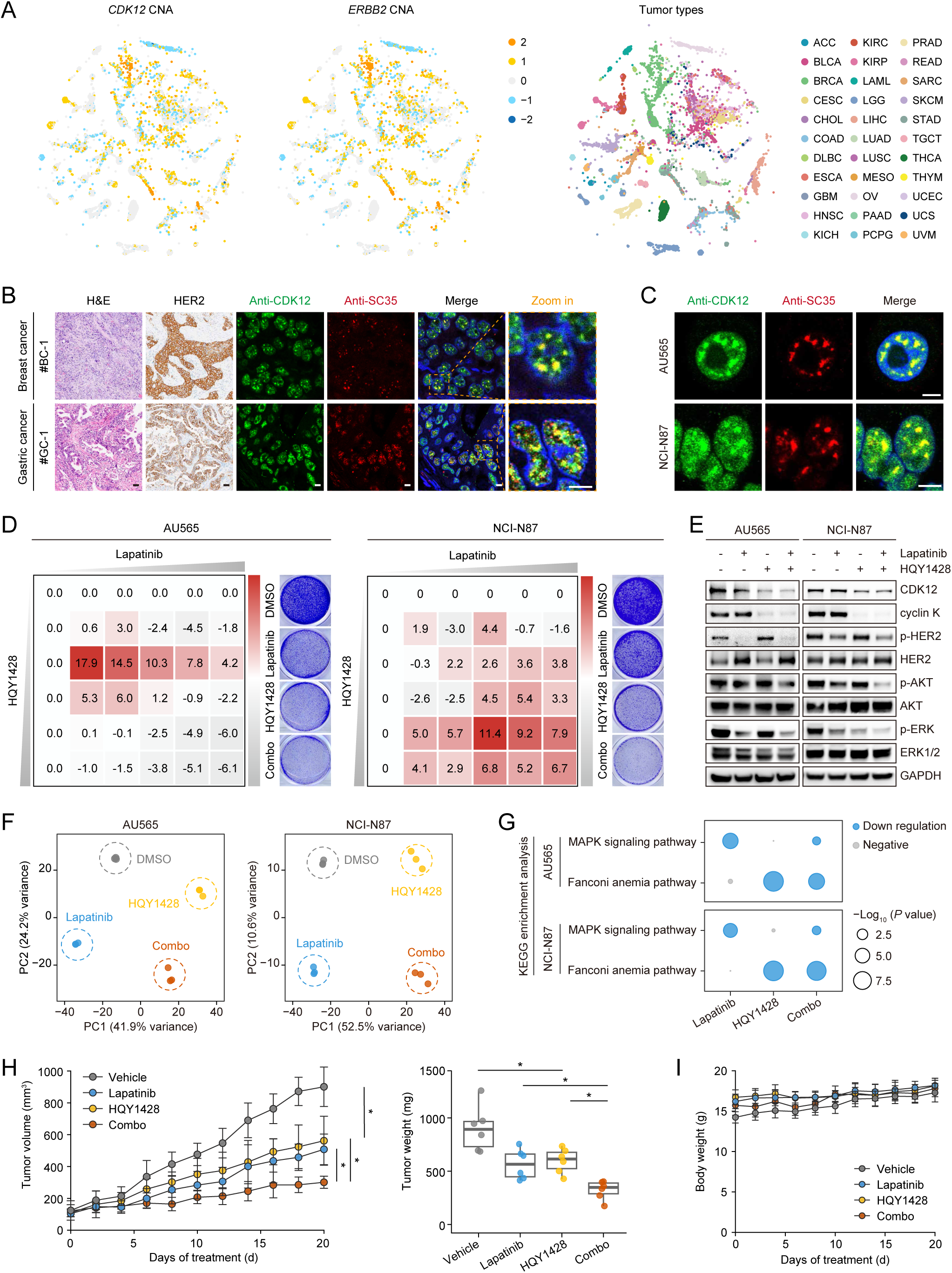
HQY1428 enhances the antitumor activity of HER2 inhibitors. A. Copy number profiles of *CDK12* and *ERBB2* across different tumor types from The Cancer Genome Atlas (TCGA) database. Each point represents a tumor sample and is colored by *CDK12* copy number status, *ERBB2* copy number status, or tumor type, as indicated. B. Representative images of hematoxylin and eosin (H&E) staining, immunohistochemistry (IHC) staining for HER2, and immunofluorescence (IF) staining for CDK12 (green) and SC35 (red) in HER2-positive breast and gastric tumor sections. Cell nuclei were counterstained with DAPI (blue). Boxed regions are enlarged on the right. Scale bar, 50 μm (H&E and IHC) or 10 μm (IF). C. Representative immunofluorescence images of CDK12 (green) and SC35 (red) in AU565 and NCI-N87 cells with *ERBB2* amplification. Cell nuclei were counterstained with DAPI (blue). Scale bar, 5 μm. D. Bliss synergy score heatmaps for lapatinib and HQY1428 combinations in AU565 and NCI-N87 cells. Representative crystal violet staining of cells treated with DMSO, lapatinib (200 nM for AU565 and 20 nM for NCI-N87), HQY1428 (4 nM for AU565 and 6 nM for NCI-N87), or their combination is shown alongside each heatmap. E. Immunoblotting analysis of the indicated proteins in AU565 and NCI-N87 cells treated with DMSO, lapatinib (200 nM for AU565 and 20 nM for NCI-N87), HQY1428 (4 nM for AU565 and 6 nM for NCI-N87), or their combination for 24 h. GAPDH was used as the loading control. F. Principal component analysis (PCA) of RNA-seq data from AU565 and NCI-N87 cells treated with DMSO, lapatinib (200 nM for AU565 and 20 nM for NCI-N87), HQY1428 (4 nM for AU565 and 6 nM for NCI-N87), or their combination for 6 h. G. KEGG enrichment analysis of the downregulated genes in AU565 and NCI-N87 cells treated with lapatinib, HQY1428, or their combination. H. Tumor growth curves (left) and tumor weight quantification (right) of NCI-N87 xenografts in BALB/c nude mice orally treated with vehicle control (0.5% hydroxypropyl methylcellulose and 0.1% Tween 80), lapatinib (30 mg/kg/day), HQY1428 (10 mg/kg/day), or their combination. Tumor growth curves are presented as mean ± s.d. (n = 6). Each point in the tumor weight plot represents one mouse. \**P* < 0.05, ANOVA followed by Tukey’s post-test. I. Body weight measurements of BALB/c nude mice during vehicle control, lapatinib, HQY1428, or combination treatment.

These findings provided a rationale for combining HER2 inhibitors with cyclin K molecular glue degraders in *ERBB2*-amplified cancers. Dose-matrix experiments demonstrated synergistic growth inhibition by lapatinib and HQY1428 in multiple *ERBB2*-amplified cell lines (Figure 7D; Supplementary Figure 9D). In AU565 and NCI-N87 cells, combination treatment resulted in reduced PI3K and MAPK signaling (Figure 7E). We then performed RNA-seq on AU565 (Supplementary Tables 12-14) and NCI-N87 cells (Supplementary Tables 15-17) treated with lapatinib, HQY1428, or both (Figure 7F). KEGG enrichment analysis identified the MAPK signaling pathway among genes downregulated by lapatinib and the Fanconi anemia pathway among genes downregulated by HQY1428 (Figure 7G). The complementary effects of combination treatment on growth signaling and DNA repair programs may contribute to the enhanced antitumor activity.

In NCI-N87 xenografts, lapatinib (30 mg/kg/day) combined with HQY1428 (10 mg/kg/day) suppressed tumor growth more effectively than either agent alone (Figure 7H). Tumor analysis showed reduced cyclin K and CDK12 levels, decreased Ki-67 staining, and increased cleaved caspase-3 staining (Supplementary Figure 10A). Importantly, there was no significant loss of body weight (Figure 7I), no overt histological abnormalities in major organs (Supplementary Figure 10B), and no remarkable changes in the measured hematological and biochemical parameters (Supplementary Figure 10C). In conclusion, these data highlight the therapeutic potential of combining HER2 inhibitors with cyclin K molecular glue degraders, offering a promising strategy for treating *ERBB2*-amplified cancers.

## Discussion

This study demonstrates that CDK12 forms dynamic liquid-like condensates at nuclear speckles, a process that can be further enhanced by CDK12 inhibitor treatment. Increased cyclin K recruitment into CDK12 condensates may contribute to reduced sensitivity to kinase inhibition. Building on these insights, we fuse a CDK12-binding module with a DDB1-engaging moiety to rationally design a new class of potent cyclin K molecular glue degraders. The lead compound HQY1428 effectively disrupts CDK12 phase separation, perturbs oncogenic transcription, and exerts potent single-agent antitumor activity, as well as synergistic efficacy with HER2 blockade in *ERBB2*-amplified malignancies (Figure 8). Our findings pinpoint a promising strategy to target CDK12 condensates through cyclin K degradation.

**Figure 8.**
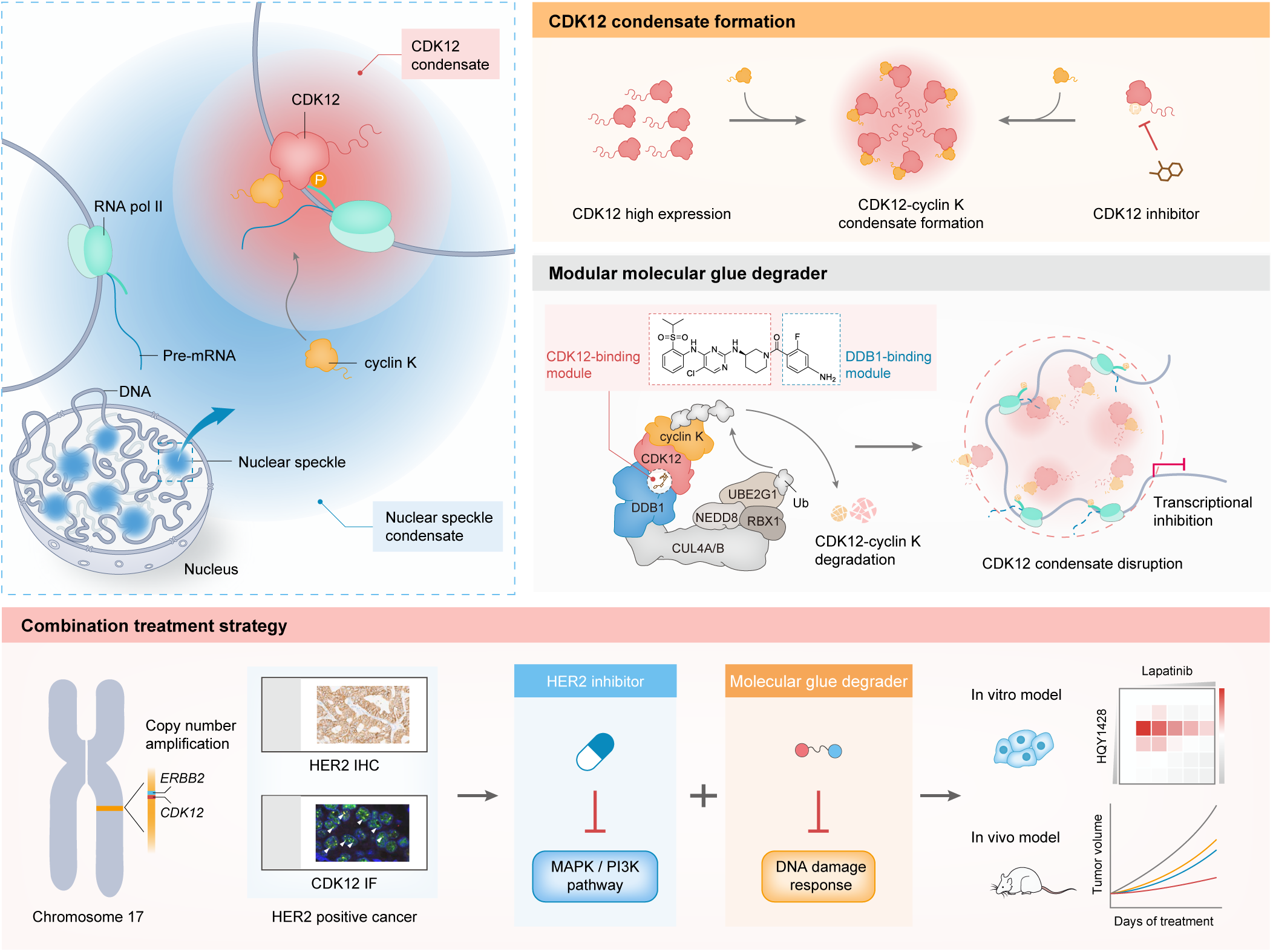
Schematic model of CDK12 condensate targeting. CDK12 forms dynamic condensates at nuclear speckles, with increased puncta accumulation linked to high CDK12 expression and CDK12 inhibitor treatment. Rationally designed cyclin K molecular glue degraders promote CDK12-DDB1 association and cyclin K degradation, accompanied by CDK12 depletion and loss of CDK12 condensates. Furthermore, *CDK12* is often co-amplified with *ERBB2* (encoding HER2) due to their genomic proximity, which provides a rationale for combining HER2 inhibitors and cyclin K molecular glue degraders to enhance antitumor activity in *ERBB2*-amplified cancers.

Previous work has established that CDK12 interacts with hyperphosphorylated RNAPII in nuclear speckles, where it phosphorylates multiple spliceosomal components within these membraneless organelles (12). CDK12 is uncommon in the CDK family to contain an arginine/serine-rich (RS) domain, which is critical for its localization (29). Here, we show that both the N- and C-terminal regions of CDK12 are extensively disordered, whereas the N-terminal IDR and RS domain are critical for condensate formation at nuclear speckles. Intriguingly, cyclin K alone did not undergo phase separation despite bearing IDR itself under our experimental conditions. Instead, CDK12 compartmentation seemed to serve as a driving force for recruiting cyclin K into nuclear speckles to assemble active CDK12-cyclin K complexes, though the precise mechanisms need to be elucidated (59). The CDK12 close homolog CDK13 may share similar properties but require further exploration because of relatively low endogenous expression. Notably, elevated CDK12 abundance resulted in increased nuclear puncta accumulation, suggesting a pivotal role of protein concentration. In addition, kinase inhibitor treatment also caused an enlargement of CDK12-cyclin K aggregates. We speculate that the π-π and cation-π interactions between small-molecules and biomolecules can modify the physicochemical features of phase-separated condensates, as proposed in recent reports (9,15,60). It remains to be determined whether myriad CDK12 inhibitors of diverse chemistry display different behaviors, and how the consequent drug partitioning would in turn influence the pharmacodynamics and activity of specific compounds.

These findings suggest that cyclin K degradation provides a means of dismantling CDK12 assemblies that persist during kinase inhibition. Recent structural studies have established principles for designing cyclin K molecular glue degraders through optimization of CDK12 binding and DDB1 engagement (61–65). We applied a modular design approach to combine these features. The resulting ZSQ253 exerted remarkable nanomolar cytotoxicity, outperforming the well-characterized cyclin K molecular glue degrader CR8. Structural optimization of ZSQ253 yielded HQY1428 with superior potency through a multi-site gluing machinery. The cryo-EM structure supported HQY1428 binding at the CDK12-DDB1 interface and suggested an additional polar contact involving its amino group. Enhanced DDB1 association was accompanied by faster cyclin K depletion, CDK12 loss, and condensate disruption. Of note, the inter-protein binding pocket for molecular glue degraders is generally hydrophobic, which often raises challenges in balancing the affinity and solubility. Nevertheless, HQY1428 showed favorable pharmacokinetic properties and in vivo antitumor efficacy. Additional pharmacological and preclinical characterization of this lead compound is underway.

Given the positive correlation between CDK12 abundance and puncta accumulation, it is appealing to employ cyclin K molecular glue degraders against human malignancies harboring *ERBB2* amplification, in which *CDK12* is frequently co-amplified. Indeed, across diverse HER2-positive tumor tissues and cell lines, we observed prominent CDK12 expression and distinct CDK12 droplets at nuclear speckles. Recent studies have uncovered that structurally coupled co-amplification of an oncogene and a passenger gene can create unique cancer dependencies and lead to collateral therapeutic vulnerabilities (66,67). Along the same line, our data indicated that combining HER2 inhibition and CDK12-cyclin K degradation achieved synergistic efficacy in *ERBB2*-amplified neoplasms without obvious adverse effects.

Although only small-molecule inhibitors were tested in the current study, other approved drug modalities such as HER2-directed antibodies (e.g., trastuzumab, pertuzumab) (68,69) and antibody-drug conjugates (e.g., trastuzumab emtansine, trastuzumab deruxtecan) (70,71) will most likely produce consistent results and warrant future investigations. Moreover, these findings provide a compelling rationale for conjugating HER2 monoclonal antibodies with cyclin K molecular glue degraders as a novel treatment option in this disease setting.

In summary, our rational design of a modular cyclin K molecular glue degrader with oral availability and antitumor potency demonstrates the feasibility and effectiveness of targeting transcriptional condensates through protein degradation. While the therapeutic potential of modulating phase separation has generated significant interest in drug discovery, directly targeting biomolecular condensates remains challenging due to the limited manipulability and specificity of intrinsically disordered regions. We offer an alternative approach by pharmacologically degrading cyclin K to disrupt CDK12 condensates, supporting further development of cyclin K molecular glue degraders, particularly in *ERBB2*-amplified cancers.

## Materials and Methods

### Cell lines and reagents

Cell lines used in this study were obtained from the American Type Culture Collection (ATCC) or Japanese Collection of Research Bioresources Cell Bank (JCRB). All cell lines were routinely tested for mycoplasma contamination, and cell identity was verified using short tandem repeat (STR) profiling. Cells were maintained in RPMI 1640 (Life Technologies) supplemented with 10% fetal bovine serum (Gibco), 2 mM L-glutamine, 1 mM sodium pyruvate, 100 U/mL penicillin, and 100 μg/mL streptomycin. Commercial small-molecule compounds were purchased from Selleck Chemicals and dissolved in DMSO (Sigma-Aldrich) at a stock concentration of 10 mM.

### Patient samples

The study was approved by the Ethics Committee of Ren Ji Hospital and written informed consent was obtained from all patients. We assembled a cohort of 22 patients diagnosed with HER2-positive cancer, as determined by immunohistochemistry. Patient characteristics are summarized in Supplementary Table 11.

### Immunofluorescence staining

For tumor cell lines, cells were grown in 8-well culture dishes (Ibidi), fixed with 4% paraformaldehyde for 15 min at room temperature (RT), and permeabilized with 0.2% Triton X-100 (Sigma-Aldrich) in PBS for 10 min. After three washes with PBS, cells were blocked with 2% bovine serum albumin (BSA) in PBS for 30 min at RT, followed by incubation with primary antibodies at 4 °C overnight. The following primary antibodies were used: CDK12 (HPA008038, Sigma-Aldrich), SC35 (ab11826, Abcam), pSer2-Pol II (ab193468, Abcam), and γH2AX (#80312, Cell Signaling Technology). After incubation, cells were washed three times with PBS and incubated with Alexa Fluor-conjugated secondary antibodies (Invitrogen) for 1 h in the dark. After three washes with PBS, cells were counterstained with 4’,6-diamidino-2-phenylindole (DAPI) (Invitrogen) for 5 min. For formalin-fixed and paraffin-embedded (FFPE) sections, slides were baked, dewaxed with xylene, rehydrated through graded alcohols, and subjected to antigen retrieval in 10 mM sodium citrate buffer (pH 6.0) using a steam pressure cooker for 20 min. Slides were then treated with 3% hydrogen peroxide (H2O2) in methanol for 10 min to quench endogenous peroxidase activity, permeabilized with 0.5% Triton X-100 in PBS for 15 min, blocked with 10% normal goat serum in PBS for 1 h at RT, and incubated with primary antibodies at 4 °C overnight. The following primary antibodies were used: CDK12 (HPA008038, Sigma-Aldrich) and SC35 (ab11826, Abcam). After three washes in PBST (PBS containing 0.1% Tween-20), slides were incubated with Alexa Fluor-conjugated secondary antibodies (Invitrogen) for 1 h in the dark. Finally, slides were washed three times in PBST, cover-slipped with mounting medium containing DAPI (Beyotime), and sealed with nail polish. Fluorescent images were acquired using a Leica TCS SP8 confocal microscopy system with a 63× oil-immersion objective or a Leica STELLARIS 5 confocal microscopy system with a 100× oil-immersion objective. Image analysis and quantification were performed using LAS X and Fiji (ImageJ) software. CDK12 puncta enrichment index was calculated as the mean intensity of the brightest 5% of nuclear pixels divided by the mean intensity of the dimmest 50% of nuclear pixels.

### Live-cell imaging and FRAP

Cells were grown in 8-well culture dishes (Ibidi) and imaged using a Leica TCS SP8 confocal microscopy system equipped with a 63× oil-immersion objective. Imaging was performed in an incubation chamber maintained at 37 °C with 5% CO₂. For the 1,6-hexanediol assay, SKOV3 cells expressing CDK12-mEGFP were imaged before and after treatment with 3% 1,6-hexanediol for 15 s. Fluorescence recovery after photobleaching (FRAP) assays were conducted using the FRAP module of the Leica TCS SP8 confocal microscopy system. CDK12-mEGFP puncta were bleached using a 488-nm laser set to 100% laser power. Time series for FRAP analysis were captured over eight cycles at 1-second intervals. Fluorescence intensity was measured in three regions of interest from separate nuclei and quantified using LAS X software.

### IncuCyte imaging

Cells were seeded at a density of 2 × 10^3^ cells per well in 96-well plates and allowed to adhere overnight before treatment with the indicated concentrations of compounds. For apoptosis kinetics, Caspase-3/7 green detection reagent (Invitrogen) was added at the time of compound treatment. Cell growth and caspase-3/7 fluorescence were monitored using the Incucyte S3 live-cell analysis system (Sartorius). Time-lapse imaging was performed at 4 or 8 h intervals. Cell confluence and green fluorescence were quantified using the ilastik software (v1.4.0).

### In vivo studies

For single-agent treatment, SKOV3 cells (3 × 10^6^) were intraperitoneally injected into female BALB/c nude mice (5 weeks of age). Upon tumor formation, mice were randomly assigned to the vehicle or treatment group (n = 6 per group). HQY1428 was administered orally at doses of 5 mg/kg/day or 10 mg/kg/day, with vehicle (2.5% v/v DMSO and 97.5% v/v 30% SBE-β-CD) as the negative control. For combination treatment, NCI-N87 cells (5 × 10^6^) were mixed with Matrigel (BD Biosciences) and injected subcutaneously into the dorsal flank of female BALB/c nude mice (5 weeks of age). When the mean tumor volume reached 120 mm^3^, mice were randomly assigned to the vehicle or treatment group (n = 6 per group). HQY1428 was administered orally at a dose of 10 mg/kg/day, while lapatinib was given orally at 30 mg/kg/day in 0.5% hydroxypropyl methylcellulose and 0.1% Tween 80. Mice were monitored and weighed every other day. For the subcutaneous model, tumor volumes were measured with a caliper and calculated as length × width^2^ × 0.5. After three weeks of treatment, mice were euthanized, and xenografts were harvested and imaged. For the intraperitoneal model, total tumor weight was calculated as the sum of the weights of all resectable implants from each mouse. Tumor tissues were fixed in 4% paraformaldehyde for immunohistochemical analysis. The heart, liver, spleen, lung, and kidney were collected for histological examination. Hematological and biochemical parameters were assessed. The Institutional Animal Care and Use Committee of Ren Ji Hospital approved the animal protocols, and all experiments were performed in accordance with Ren Ji Hospital policies regarding the care, welfare, and treatment of laboratory animals.

## Statistical analysis

The RNA-seq and copy number datasets for 33 TCGA cancer types were obtained from the Pan-Cancer Atlas and downloaded from the UCSC Xena Explorer (cohort: TCGA Pan-Cancer). The RNA-seq and copy number datasets for cancer cell lines were retrieved from the Cancer Cell Line Encyclopedia (CCLE). Pearson’s correlation coefficient was used to measure the linear correlation between two variables. For continuous variables, differences between two groups were assessed using the two-sided Student’s *t*-test, whereas one-way or two-way analysis of variance (ANOVA) was used for comparisons among multiple groups or experimental factors. Tukey’s post-test was used for the ANOVA comparisons. The Kruskal-Wallis test was used for nonparametric comparisons among multiple groups. Graphs were generated and statistical analyses were performed with GraphPad Prism (v9.0) or R (v4.1.0). *P* values < 0.05 were considered statistically significant.

## Data availability

The sequencing data were deposited in the NCBI BioProject database under the accession number PRJNA1238237.

## Acknowledgements

This work was supported by the National Natural Science Foundation of China (82373351 and 82573385 to G Zhuang; 82674304 to P Shen; 82605747 to K Ye; 82403676 to L Cheng; 82373972 to D Zhao; 82473277 to F Yang), the Strategic Priority Research Program of the Chinese Academy of Sciences Grant (XDB1060000 to L Tan, K Ding, and Z Wang; XDB0830300 to D Zhao), Shanghai Pujiang Program (23PJ1407600 to F Yang), Jilin Provincial Health Talent Special Program (2024WSZX-D04 to L Cheng), Shanghai Basic Research Pioneer Project (L Tan, Y-X Zhang, and Y-Y Zhang), Shanghai Municipal Science and Technology Major Project (L Tan, Y-X Zhang, and Y-Y Zhang), Shanghai Science and Technology Commission Project (23JC1403000 to W Di; 21S11900300 to L Xiang), Chongqing Natural Science Foundation Postdoctoral Fund (cstc2021jcyj-bshX0048 to D Chen), the General Program of Chongqing Natural Science Foundation (cstc2021jcyj-msxmX0037 to Z-L Wang), Guizhou Provincial Science and Technology Achievements Application and Industrialization Plan (LC [2021]012 to X Wang), and Guizhou Provincial Science and Technology Projects (grant number Qiankeherencai XKBF [2025]024 to G Zhuang).

