## Supplementary Method for "Targeting CDK12 condensates at nuclear speckles with rationally designed cyclin K molecular glue degraders"

### Supplementary Methods

#### Prediction of intrinsic disorder and protein structure

Intrinsically disordered regions (IDRs) in CDK12 and cyclin K were predicted using PONDR VSL2 (https://www.pondr.com/) and IUPred3 in long-disorder mode (https://iupred3.elte.hu/) (1,2). Residues with disorder scores above 0.5 were classified as predicted disordered residues. Predicted three-dimensional structures of both proteins were retrieved from the AlphaFold Protein Structure Database (https://alphafold.ebi.ac.uk/), which provides protein structure models generated using AlphaFold2 (3,4).

#### Cell viability and drug combination assays

Cell viability was assessed using crystal violet staining or the Cell Counting Kit-8 (CCK-8) assay (ShareBio). For crystal violet staining, cells were seeded at $2 \times 10^4$ cells per well in 12-well plates, allowed to adhere overnight, and treated with serial dilutions of the indicated compounds for 5 days. Cells were then fixed with formalin and stained with 0.1% crystal violet (Sigma-Aldrich). Staining was quantified by densitometric analysis. For the CCK-8 assay, cells were seeded in triplicate in 96-well plates at $3 \times 10^3$ cells per well and subjected to the indicated treatments for 72 h before measuring absorbance at 450 nm. To evaluate drug combination effects, the expected fractional growth inhibition under the Bliss independence model was calculated as E(Bliss) = A + B − A × B. A and B represent the fractional growth inhibition induced by each agent alone at the corresponding dose. Bliss excess was calculated as E(observed) − E(Bliss), where E(observed) denotes the fractional growth inhibition produced by the combination. Positive Bliss excess values indicate greater growth inhibition than expected under the Bliss independence model. Bliss synergy scores were calculated as 100 × [E(observed) − E(Bliss)].

**Plasmids and sgRNAs**

The *CDK12* and *DDB1* open reading frames (ORFs) were amplified from the cDNA of HEK293T cells, while the *CCNK* ORF was synthesized by GenScript. Plasmids expressing full-length CDK12, CDK12 deletion mutants, cyclin K, and DDB1 were constructed using the ClonExpress Ultra One Step Cloning Kit (Vazyme) by homology-based assembly directly into pLenti7.3/V5-DEST. The CRISPR-Cas9 system was employed to knock out the indicated genes. The primers for cloning and sgRNA sequences are provided in Supplementary Table 18 and Supplementary Table 19, respectively.

**Virus production and cell infection**

HEK293T cells in a 10-cm dish were transfected with 6 μg of LentiCRISPRv2 vector encoding the indicated sgRNA, 13.8 μg of plasmid Δ8.9, and 1.2 μg of plasmid VSV-G using Lipofectamine 2000 transfection reagent (Invitrogen). Cells were incubated at 37 °C and the medium was replaced after 12 h. Viral supernatants were collected at 72 h after transfection and supplemented with 8 μg/mL polybrene (Fluka) to infect SKOV3 cells seeded in 6-well dishes. Infected cells were selected with 3 μg/mL puromycin (Selleck Chemicals) for one week to establish stable cell lines. Knockout efficiency was assessed by immunoblotting for the corresponding proteins.

**Western blot analysis**

Cells were lysed in RIPA buffer (50 mM Tris pH 7.4, 150 mM NaCl, 1% NP-40, 0.1% SDS, 2 mM EDTA) containing protease inhibitors (Roche) and phosphatase inhibitors (Roche). Protein concentrations were quantified using Pierce BCA Protein Assay Kit (Thermo Fisher Scientific). Whole-cell lysates (~20 μg protein) were separated on NuPAGE 10% Bis-Tris gels (Invitrogen) and transferred onto nitrocellulose membranes. Membranes were blocked with 5% non-fat milk in TBST (20 mM Tris pH 7.4, 150 mM NaCl, 0.05% Tween-20) for 1 h and incubated with primary antibodies at 4 °C overnight, followed by incubation with horseradish peroxidase (HRP)-conjugated secondary antibodies (Cell Signaling Technology). Protein bands were visualized by chemiluminescence with a ChemiDoc XRS+ system (Bio-Rad). The following primary antibodies were used: CDK12 (#11973, Cell Signaling Technology), cyclin K (sc-376371, Santa Cruz), CDK7 (#2916, Cell Signaling Technology), CDK9 (#2316, Cell Signaling Technology), DDB1 (#6998, Cell Signaling Technology), RBX1 (#11922, Cell Signaling Technology), CUL4A

(#2999, Cell Signaling Technology), CUL4B (ab227724, Abcam), pSer2-Pol II (ab193468, Abcam), RAD51 (#11255-1-AP, Proteintech), γH2AX (#9718, Cell Signaling Technology), p-HER2 (#2243, Cell Signaling Technology), HER2 (#4290, Cell Signaling Technology), PathScan Multiplex Western Cocktail I (#5301, Cell Signaling Technology), GAPDH (#60004-1-Ig, Proteintech), and β-actin (#5125, Cell Signaling Technology).

**Compound synthesis**

Reagents and solvents were purchased from commercial sources and used without further purification. $^{1}$H NMR spectra were recorded on a Bruker spectrometer operating at 400 MHz using tetramethylsilane (TMS) as an internal standard. Chemical shifts (δ) are reported in ppm and coupling constants (J) are reported in Hz. The signal multiplicities are denoted as follows: s, singlet; d, doublet; t, triplet; q, quartet; dd, doublet of doublets; dt, doublet of triplets; ddd, doublet of doublets of doublets; br, broad; brs, broad singlet; m, multiplet. Mass spectra were acquired using an Agilent 6120 quadrupole LC/MS system equipped with an electrospray ionization (ESI) source. High-performance liquid chromatography (HPLC) purification was performed with a Waters system equipped with a 2489 UV/Vis detector using a SunFire® Prep C18 OBD™ 5 μm column (19 mm × 100 mm) eluted with $CH_3CN/H_2O$ (0.05% TFA) at a flow rate of 15.0 mL/min with UV detection at 214/254/280 nm. The purity of all tested compounds was >95%, as determined by

HPLC and $^{1}$H NMR analyses. Silica gel column chromatography was performed with a Teledyne ISCO medium pressure system using prepackaged columns. The following abbreviations were used in this section: DIEA = *N*,*N*-diisopropylethylamine; NMP = 1-methyl-2-pyrrolidinone; DCM = dichloromethane; TFA = trifluoroacetic acid; HATU = 2-(7-azabenzotriazol-1-yl)-*N*,*N*,*N'*,*N'*-tetramethyluronium hexafluorophosphate; EA = ethyl acetate; RT = room temperature.

The synthetic routes to intermediates **2** and **3** and the final compounds are summarized in Supplementary Figure 2B.

Reagents and conditions. (a) *tert*-butyl (*R*)-3-aminopiperidine-1-carboxylate, DIEA, NMP, 135 °C, 18 h; (b) TFA, DCM, RT, 8 h; (c) the corresponding carboxylic acids, HATU, DIEA, DMSO, RT, 0.5 h.

***tert*-butyl (*R*)-3-((5-chloro-4-((2-(isopropylsulfonyl)phenyl)amino)pyrimidin-2-yl)amino)piperidine-1-carboxylate (2)**

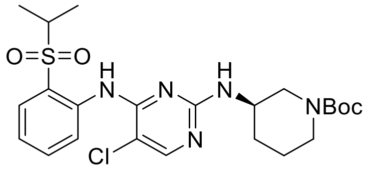

A solution of 2,5-dichloro-*N*-(2-(isopropylsulfonyl)phenyl)pyrimidin-4-amine (1.73 g, 5.00 mmol), *tert*-butyl (*R*)-3-aminopiperidine-1-carboxylate (1.00 g, 5.00 mmol), and DIEA (2.48 mL, 15.00 mmol) in NMP (20 mL) was heated at 135 °C for 18 h. After cooling to room temperature, the reaction mixture was diluted with EA (200 mL) and washed with $H_2O$ (50 mL × 3). The organic layer was dried over $Na_2SO_4$, filtered and concentrated in a rotary evaporator. The crude residue was then purified by C18 flash column chromatography ($H_2O$/MeCN = 100:0 to 20:80). The product, **2**, was obtained by vacuum drying as yellow solid (2.32 g, 4.56 mmol) with a yield of 91.20%. MS-ESI: m/z calculated for $C_{23}H_{32}ClN_5O_4S$, Exact Mass: 509.19, found 510.14 $[M + H]^+$.

**(*R*)-5-chloro-*N*$^4$-(2-(isopropylsulfonyl)phenyl)-*N*$^2$-(piperidin-3-yl)pyrimidine-2,4-diamine (3)**

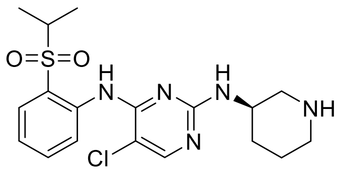


A solution of **2** (2.32 g, 4.56 mmol) and TFA (5 mL) in DCM (50 mL) was stirred at room temperature for 8 h. DCM was removed by a rotary evaporator, and the crude residue was purified by C18 flash column chromatography ($H_2O$/MeCN = 100:0 to 40:60). The product, **3**, was obtained by vacuum drying as white solid (1.62 g, 3.96 mmol) with a yield of 86.86%. MS-ESI: m/z calculated for $C_{18}H_{24}ClN_5O_2S$, Exact

Mass: 409.13, found 410.09 [M + H]$^+$. $^1$H NMR (400 MHz, $CDCl_3$) δ 9.53 (s, 1H), 8.67 (d, *J* = 8.4 Hz, 1H), 8.02 (s, 1H), 7.88 (dd, *J* = 7.9, 1.1 Hz, 1H), 7.68 – 7.59 (m, 1H), 7.20 (t, *J* = 7.6 Hz, 1H), 5.39 (s, 1H), 5.30 (s, 1H), 3.89 (s, 1H), 3.33 – 3.21 (m, 1H), 3.20 – 3.13 (m, 1H), 2.94 – 2.85 (m, 1H), 2.81 – 2.72 (m, 1H), 2.72 – 2.64 (m, 1H), 2.03 – 1.87 (m, 1H), 1.86 – 1.72 (m, 1H), 1.64 – 1.48 (m, 2H), 1.31 (dd, *J* = 6.8, 2.8 Hz, 6H).

**(*R*)-(3-((5-chloro-4-((2-(isopropylsulfonyl)phenyl)amino)pyrimidin-2-yl)amino)piperidin-1-yl)(phenyl)methanone (ZSQ253)**

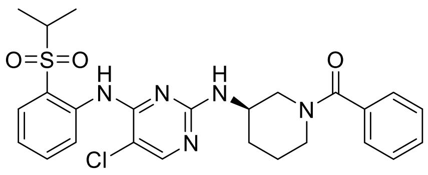

A solution of **3** (409 mg, 1.00 mmol), benzoic acid (122 mg, 1.00 mmol), HATU (380 mg, 1.00 mmol), and DIEA (198 μL, 1.20 mmol) in DMSO (5 mL) was stirred at room temperature for 0.5 h, and then the reaction mixture was purified by high-performance liquid chromatography ($H_2O$/MeCN = 90:10 to 25:75). The product, **ZSQ253**, was obtained by vacuum drying as white solid (262 mg, 0.511 mmol) with a yield of 51.07%. MS-ESI: m/z calculated for $C_{25}H_{28}ClN_5O_3S$, Exact Mass: 513.16, found 514.20 [M + H]$^+$. $^1$H NMR (400 MHz, $CDCl_3$) δ 10.46 (s, 1H), 10.11 (s, 1H), 8.59 (s, 1H), 7.96 (d, *J* = 7.6 Hz, 1H), 7.86 – 7.76 (m, 2H), 7.54 – 7.34 (m, 6H), 4.54 – 4.39 (m, 1H), 4.11 – 3.94 (m, 1H), 3.65 – 3.48 (m, 1H), 3.34 – 3.26 (m, 1H), 3.25 –

3.15 (m, 2H), 2.17 – 2.04 (m, 1H), 1.94 – 1.81 (m, 2H), 1.61 – 1.46 (m, 1H), 1.35 (d, *J* = 6.4 Hz, 3H), 1.31 (d, *J* = 6.7 Hz, 3H).

**(*R*)-(3-amino-5-fluorophenyl)(3-((5-chloro-4-((2-(isopropylsulfonyl)phenyl)amino)pyrimidin-2-yl)amino)piperidin-1-yl)methanone (HQY1428)**

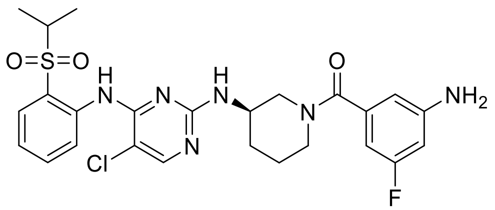


A solution of **3** (409 mg, 1.00 mmol), 3-amino-5-fluorobenzoic acid (155 mg, 1.00 mmol), HATU (380 mg, 1.00 mmol), and DIEA (198 μL, 1.20 mmol) in DMSO (5 mL) was stirred at room temperature for 0.5 h, and then the reaction mixture was purified by high-performance liquid chromatography ($H_2O$/MeCN = 90:10 to 35:65). The product, **HQY1428**, was obtained by vacuum drying as white solid (215 mg, 0.394 mmol) with a yield of 39.38%. MS-ESI: m/z calculated for $C_{25}H_{28}ClFN_6O_3S$, Exact Mass: 546.16, found 547.18 $[M + H]^+$. $^1H$ NMR (400 MHz, $CDCl_3$) δ 10.36 (s, 1H), 9.10 (s, 1H), 8.54 – 8.22 (m, 2H), 7.92 (d, *J* = 7.6 Hz, 1H), 7.88 – 7.79 (m, 1H), 7.79 – 7.70 (m, 1H), 7.66 – 7.54 (m, 1H), 7.43 (t, *J* = 6.3 Hz, 1H), 7.26 – 7.05 (m, 1H), 4.34 – 3.80 (m, 2H), 3.75 – 3.49 (m, 2H), 3.47 – 3.13 (m, 2H), 2.14 – 1.70 (m, 3H), 1.61 – 1.52 (m, 1H), 1.28 (s, 6H).

**(*R*)-1-(3-((5-chloro-4-((2-(isopropylsulfonyl)phenyl)amino)pyrimidin-2-yl)amino)piperidin-1-yl)ethan-1-one (ZSQ2549)**

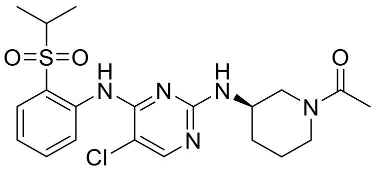


A solution of **3** (40.9 mg, 0.10 mmol), acetic acid (6 mg, 0.10 mmol), HATU (38 mg, 0.10 mmol), and DIEA (20 μL, 0.12 mmol) in DMSO (0.5 mL) was stirred at room temperature for 0.5 h, and then the reaction mixture was purified by high-performance liquid chromatography ($H_2O$/MeCN = 90:10 to 25:75). The product, **ZSQ2549**, was obtained by vacuum drying as white solid (33.1 mg, 0.073 mmol) with a yield of 73.39%. MS-ESI: m/z calculated for $C_{20}H_{26}ClN_5O_3S$, Exact Mass: 451.14, found 452.16 $[M + H]^+$. $^1H$ NMR (400 MHz, $CDCl_3$) $\delta$ 9.56 (s, 1H), 8.55 (d, *J* = 8.3 Hz, 1H), 8.05 (s, 1H), 7.89 (t, *J* = 7.9 Hz, 1H), 7.70 – 7.58 (m, 1H), 7.26 – 7.17 (m, 1H), 5.03 (s, 1H), 3.98 – 3.80 (m, 2H), 3.30 – 3.06 (m, 3H), 2.10 – 2.02 (m, 3H), 1.93 – 1.73 (m, 3H), 1.65 – 1.57 (m, 2H), 1.33 – 1.27 (m, 6H).

**Induced-fit docking**

Induced-fit docking was performed using the Induced Fit Docking (IFD) module of the Schrödinger Suite 2021. The protein structure of the DDB1-CR8-CDK12-cyclin K complex (PDB ID: 6TD3) was retrieved from the Protein Data Bank and prepared using the Protein Preparation Wizard in Maestro. For the compounds ZSQ253 and

HQY1428, the LigPrep module of the Schrödinger Suite was utilized to generate ionization states, tautomeric states, and stereoisomers. The standard IFD protocol was followed, with default receptor and ligand van der Waals scaling factors of 0.50 to allow flexibility during ligand docking. Twenty poses were retained from the initial docking step. Residues within 5.0 Å of the ligand poses, including side chains, were further refined using Prime to relax the protein structure and accommodate ligand-induced conformational changes. Subsequently, ligands were redocked using Glide standard precision (SP) into receptor structures with Prime energies within 30.0 kcal/mol of the lowest-energy refined structure. The resulting IFD poses were evaluated and selected for subsequent molecular dynamics simulations.

**Molecular dynamics simulations**

Molecular dynamics simulations were performed for CDK12-cyclin K-DDB1 complexes bound to CR8, ZSQ253, or HQY1428 using the Desmond module of the Schrödinger suite. The complexes were solvated with the TIP3P water model in an orthorhombic solvation box, including a 10 Å buffer size between the solutes and the edges of the box. The systems were neutralized with 28 $Na^+$ counterions, and NaCl was added to a concentration of 0.15 M. The OPLS4 force field was applied to the simulation systems. The systems were then equilibrated using a relaxation protocol prior to the production runs. Molecular dynamics (MD) simulations were performed for 200 ns for each protein-ligand complex under an NPT ensemble at 300 K and

1.01325 bar. Data were recorded at 200-ps intervals with an integration time step of 2 fs, resulting in 1000 recorded frames per simulation. Post-MD analyses were conducted using simulation interaction protocols to evaluate protein-ligand interactions.

**Cryo-EM sample preparation and data acquisition**

Protein sample quality was initially assessed by negative-stain electron microscopy using a Tecnai transmission electron microscope operated at 120 kV. Protein concentrations were measured using a NanoPhotometer spectrophotometer (IMPLEN). Purified samples (4 µL) were applied to glow-discharged 200-mesh Quantifoil 2/1 gold grids. After a 5-second incubation, the grids were blotted for 2 seconds and plunge-frozen in liquid ethane using an FEI Vitrobot Mark IV, operated at 4 °C and 100% humidity. The grids were transferred to a Titan Krios G4 transmission electron microscope operating at 300 kV, equipped with a BioContinuum K3 direct electron detector and a GIF energy filter (Gatan). Automated data collection was performed using EPU at a nominal magnification of 81,000×, corresponding to a calibrated pixel size of 1.055 Å. The dose rate was set to 25 electrons per pixel per second, and movies were recorded with 40 frames over a total exposure time of 2.2 seconds. Data were collected with a defocus range of −1.4 to −2.4 µm.

### Cryo-EM image processing

For the CDK12-cyclin K-DDB1-HQY1428 complex dataset, a total of 5,001 movies were collected in two independent data acquisition sessions. Movie frames were motion-corrected and dose-weighted, and contrast transfer function (CTF) parameters were estimated using CryoSPARC. Following quality assessment, 3,981 micrographs were selected for subsequent processing. An initial set of 1,032,312 particles was picked using Gautomatch without templates and extracted into 240 × 240 pixel boxes. The extracted particles were subjected to 2D classification, and classes displaying clear views of the complex were selected as templates for a second round of particle picking using Gautomatch, yielding 1,448,481 particles. These particles were extracted and subjected to 2D classification, from which 865,694 particles belonging to classes with well-defined structural features were selected for further processing in CryoSPARC. Two parallel rounds of ab initio reconstruction were performed, and particles from classes displaying clear structural features were selected for subsequent rounds of ab initio reconstruction. Through iterative classification and particle selection, poorly resolved particles were progressively excluded, yielding a final set of 192,202 particles. These particles were subjected to non-uniform (NU) refinement with C1 symmetry, resulting in a final density map at an overall resolution of 3.76 Å.

### Cryo-EM model building and structure refinement

The structure of DDB1 bound to DS17-engaged CDK12-cyclin K (PDB: 8BU1) was used as the starting model. The CDK12-cyclin K and DDB1 models were fitted into the cryo-EM density map based on the corresponding density features. A difference map was then calculated to identify additional density not accounted for by the protein models. Based on the shape and features of the unassigned density, compound HQY1428 was fitted into the corresponding region. The protein and ligand models were further refined through iterative cycles of real-space refinement in PHENIX and manual adjustment in Coot, resulting in the final CDK12-cyclin K-DDB1-HQY1428 complex model.

**Co-immunoprecipitation (co-IP) assay**

HEK293T cells were transfected with empty vector (*EV*) or FLAG-tagged DDB1. Cells were pre-treated with 0.5 μM MLN4924 for 2 h, followed by treatment with DMSO, HQY1428 (0.1 *μ* M), ZSQ253 (0.1 μM), or CR8 (0.1 μM) for 1 h. Cells were lysed in IP lysis buffer (50 mM Tris-HCl pH 8.0, 150 mM NaCl, 1% NP-40, 5% glycerol, 1 mM EDTA) supplemented with protease inhibitor cocktail (Roche). Cell lysates were centrifuged at 16,000 × g for 20 min at 4 °C to remove cell debris and then FLAG-DDB1 was immunoprecipitated with anti-FLAG M2 beads (Sigma-Aldrich) at 4 °C overnight. The beads were washed five times with IP lysis buffer and the precipitated protein complexes were eluted by heating in SDS-PAGE sample buffer for 10 min at 95 °C. The supernatants were collected and analyzed by immunoblotting.

**Mass spectrometry**

SKOV3 cells were seeded at a density of $2 \times 10^6$ cells in a 10-cm dish 24 h before treatment. Cells were treated in triplicate with 0.1 µM HQY1428 or ZSQ2549. After 4 h of incubation, cells were harvested and lysed in 1% SDS lysis buffer (1% SDS, 50 mM Tris-HCl pH 7.4, 150 mM NaCl, 10% glycerol, 10 mM β-glycerol phosphate) containing protease inhibitors (Roche). Protein concentrations were quantified using Pierce BCA Protein Assay Kit (Thermo Fisher Scientific). Cell lysates were precipitated in cold acetone at –20 °C overnight. The pellet was air-dried and resuspended in urea buffer (8 M urea, 100 mM Tris, pH 8.5), followed by reduction, alkylation, and digestion with trypsin at a 1:50 enzyme-to-protein ratio at 37 °C overnight. The digested peptides were desalted using C18-tips for subsequent liquid chromatography-mass spectrometry (LC-MS/MS) analysis. For LC-MS/MS analysis, as described previously (5), an on-line EASY-nLC 1000 HPLC coupled with an Orbitrap Fusion mass spectrometer (Thermo Fisher Scientific) was used. The peptide mixtures were directly loaded onto a 15-cm homemade capillary column (100 µm I.D., C18-AQ 1.9 µm resin, Dr. Maisch GmbH) and separated by a gradient at a flow rate of 300 nL/min. Mobile phase A consisted of 0.1% formic acid, 2% acetonitrile and 98% $H_2O$, and mobile phase B consisted of 0.1% formic acid, 2% $H_2O$ and 98%

acetonitrile. Mass spectra were acquired in a data-dependent mode with one full scan (m/z: 350-1500; resolution: 15,000; AGC target value: 3,000,000 and maximal injection time: 20 ms), followed by MS2 scan (32% normalized collision energy; AGC target value: 100,000; maximal injection time: Dynamic). MS/MS raw spectra were processed using MaxQuant software (version 1.6.0.1) against the SwissProt human protein sequences database. Trypsin was set as the digestion enzyme, and a maximum of two missed cleavages was allowed. The precursor mass tolerance and the fragment mass tolerance were set to 20 ppm and 0.1 Da, respectively. The false discovery rates at the peptide spectral match level and the protein level were both controlled below 1%. The intensities were used for protein quantification. Proteins without missing values across all three replicates in each group were retained for further analysis. Proteins with adjusted *P* values < 0.05 and $|\log_2 \text{fold change}| > 1$ were considered significant.

**EdU incorporation assay**

The EdU (5-ethynyl-2'-deoxyuridine) incorporation assay was performed with the Cell-Light™ EdU DNA Cell Proliferation Kit (RiboBio) according to the manufacturer's instructions. Briefly, after treatment with the indicated compounds for 24 h, cells were incubated in complete medium containing 50 μM EdU for 2 h. Following incubation, cells were fixed with 4% paraformaldehyde for 30 min at RT, stained with ApolloGreen fluorescent dye, and subsequently counterstained with

DAPI. Images were acquired using a Leica TCS SP8 confocal microscopy system with a 40× water-immersion objective.

**Flow cytometry**

For cell cycle analysis, cells were treated with the indicated compounds for 48 h. After trypsinization, $2 \times 10^5$ cells were fixed with 70% ethanol at –20 °C overnight, incubated with RNase A (ShareBio), and subsequently stained with propidium iodide (PI) for 15 min. Cell cycle distribution was analyzed using a flow cytometer (BD Biosciences). For apoptosis analysis, cells were treated with the indicated compounds for 72 h. Following EDTA-free trypsinization, $2 \times 10^5$ cells were collected by centrifugation at 300 × g for 5 min and washed with cold PBS, followed by staining with Annexin V-FITC and PI (ShareBio). Apoptosis was analyzed using a flow cytometer (BD Biosciences). The data were processed with the FlowJo software.

**RNA sequencing and analysis**

Cells were treated with the indicated compounds for 6 h. Each experimental condition was performed in triplicate, and total RNA was purified using the RNeasy Plus Mini Kit (Qiagen) according to the manufacturer's instructions. RNA quality was assessed using a NanoDrop 8000 (Thermo Fisher Scientific) and agarose gel electrophoresis. A total amount of 1 μg RNA was used for library preparation. Sequencing libraries were prepared using the NEBNext Ultra RNA Library Prep Kit for Illumina (NEB).

The libraries were sequenced on an Illumina NovaSeq platform to generate 30 million 150-bp paired-end reads per sample (Novogene). Downstream analyses were based on the clean data which were obtained from FASTQ raw files by removing adapter, low-quality reads, and poly-N sequences. The index of the reference genome was built using Bowtie (6), and paired-end clean reads were aligned to the human reference genome using HISAT2 (7). FeatureCounts was used to count the number of reads mapped to each gene (8). Fragments per kilobase of transcript per million mapped reads (FPKM) for each gene was calculated based on the length of the gene and the number of reads mapped to each gene. Differential expression analysis was performed on raw gene counts using the edgeR package (v4.2.1) (9). Genes with adjusted $P$ values $< 0.05$ and $|\log_2 \text{fold change}| > 1$ were considered differentially expressed. Gene set enrichment analysis was performed using the GSEA software (v4.1.0). Gene Ontology and pathway enrichment analyses were performed using Metascape, and enrichment network visualizations were generated with Cytoscape. Alternative splicing was analyzed using rMATS (v4.1.1) (10). Alternative splicing events were considered significant if FDR $< 0.05$ and $|\Delta PSI| \geq 0.1$. Intronic polyadenylation (IPA) was analyzed by IPAFinder (11). IPA events with adjusted $P < 0.05$ and |IPUI difference| > 0.1 were considered significant.

**Immunohistochemistry**

Formalin-fixed, paraffin-embedded (FFPE) tissue sections were baked, dewaxed with xylene, rehydrated through graded alcohols, and subjected to antigen retrieval in 10 mM sodium citrate (pH 6.0) using a steam pressure cooker for 20 min. The slides were then treated with 3% $H_2O_2$ in methanol for 10 min to quench endogenous peroxidase activity, blocked with 10% normal goat serum in PBS for 1 h, and incubated overnight at 4 °C with primary antibodies against CDK12 (HPA008038, Sigma-Aldrich), cyclin K (ab85854, Abcam), Ki-67 (#9027, Cell Signaling Technology), cleaved caspase-3 (#9661, Cell Signaling Technology), cleaved PARP (#5625, Cell Signaling Technology), and γH2AX (#80312, Cell Signaling Technology), followed by incubation with horseradish peroxidase-conjugated secondary antibodies for 1 h at RT. Antigen visualization was performed using 3,3'-diaminobenzidine (DAB) chromogen (Servicebio). The slides were counterstained with hematoxylin, dehydrated, and coverslipped with mounting solution (Invitrogen). Whole slides were scanned with an Aperio ScanScope system (Leica Biosystems).
