## Supplementary Figure for "Targeting CDK12 condensates at nuclear speckles with rationally designed cyclin K molecular glue degraders"

Supplementary Figure 1.

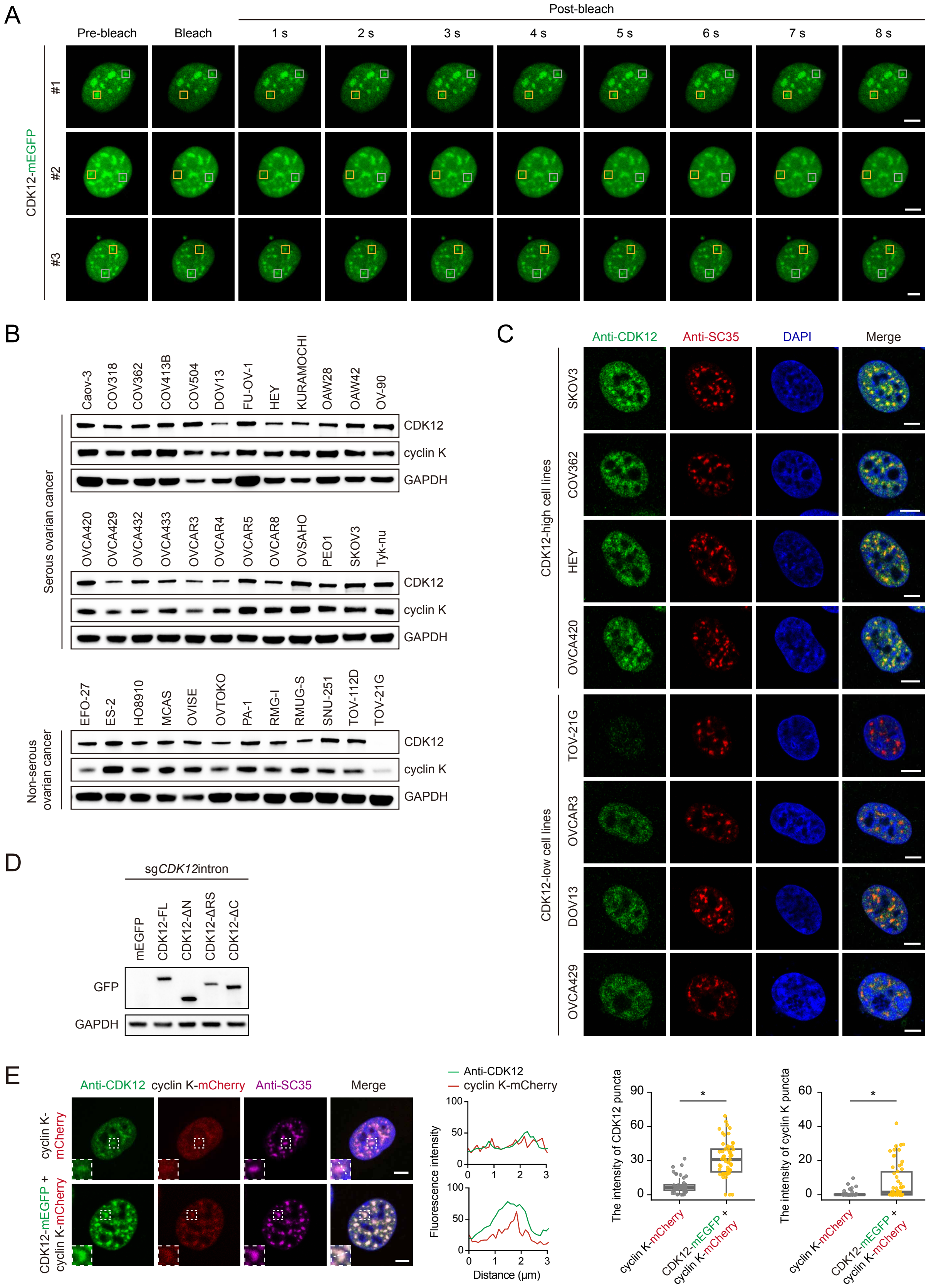


**Supplementary Figure 1.**
**A.** Time-lapse imaging of fluorescence recovery after photobleaching (FRAP) in SKOV3 cells overexpressing CDK12-mEGFP. Times relative to photobleaching are indicated. Scale bars, 5 μm. **B.** Immunoblotting analysis of CDK12 and cyclin K in a panel of ovarian cancer cell lines. GAPDH was used as the loading control. **C.** Representative immunofluorescence images of CDK12 (green) and SC35 (red) in the indicated ovarian cancer cell lines. Cell nuclei were counterstained with DAPI (blue). Scale bars, 5 μm. **D.** Immunoblotting analysis of mEGFP and the indicated mEGFP-tagged CDK12 constructs expressed in SKOV3 cells with endogenous CDK12 knockout. GAPDH was used as the loading control. **E.** Representative immunofluorescence images of CDK12 (green), cyclin K-mCherry (red), and SC35 (magenta) in SKOV3 cells overexpressing cyclin K-mCherry alone or co-expressing CDK12-mEGFP and cyclin K-mCherry. Cell nuclei were counterstained with DAPI (blue). Boxed regions are enlarged in the insets. Fluorescence intensity profiles along the indicated lines are shown in the middle. Box plots quantifying the fluorescence intensities of CDK12 and cyclin K puncta are shown on the right. Each point represents a single cell. *$P < 0.05$, two-sided Student's t-test. Scale bars, 5 μm.

Supplementary Figure 2.

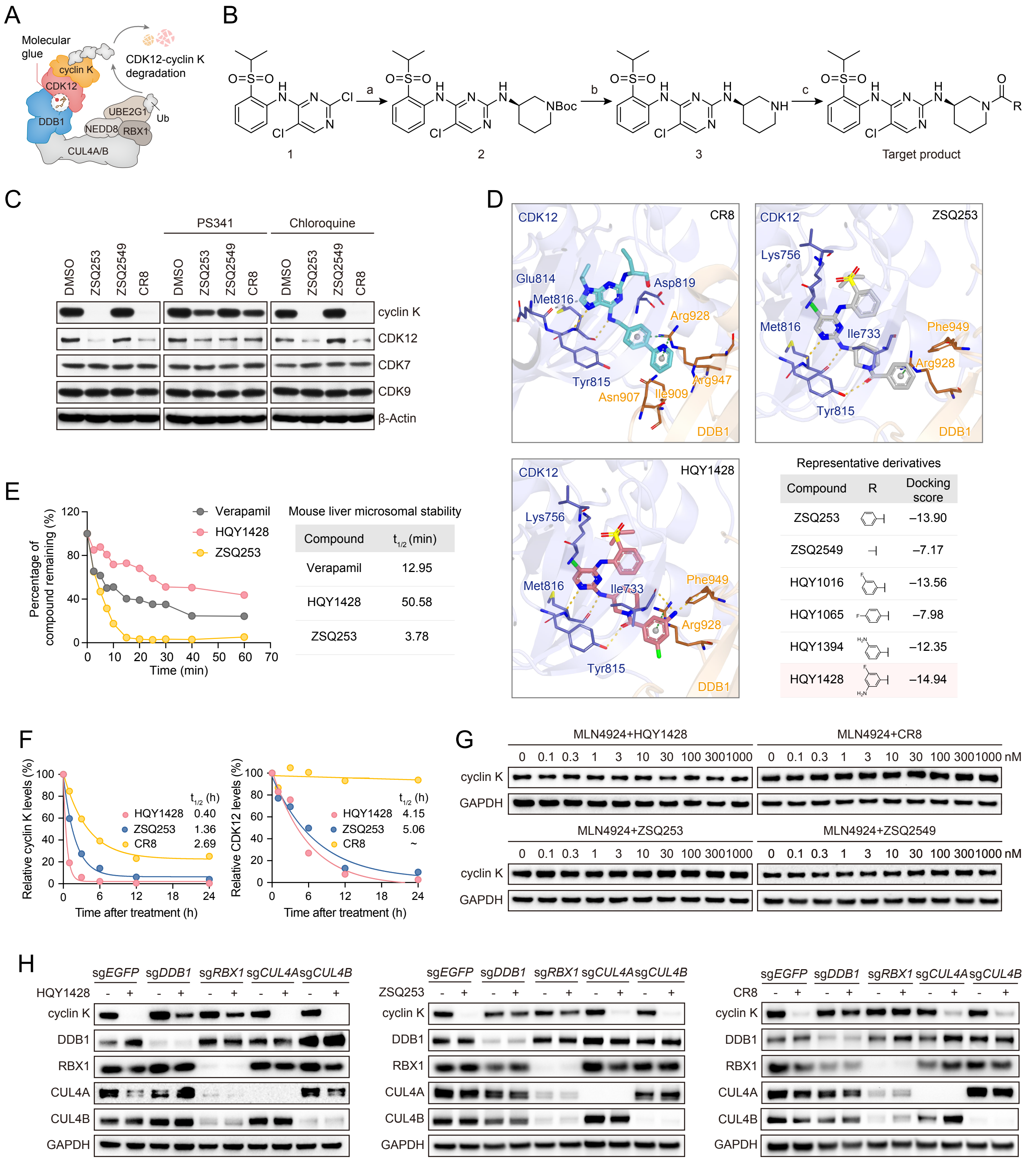


**Supplementary Figure 2.**

**A.** Schematic model of the proposed recruitment of the CDK12-cyclin K complex to the DDB1-CUL4-RBX1 E3 ubiquitin ligase machinery by cyclin K molecular glue degraders, leading to cyclin K ubiquitination and degradation, accompanied by CDK12 depletion. **B.** General synthetic route for ZSQ253, ZSQ2549, and HQY1428. The terminal substituent R distinguishes the three compounds. **C.** Immunoblotting analysis of cyclin K and CDK12 degradation in cancer cells that were pretreated with PS341 (0.1 μM) or chloroquine (10 μM) for 2 h, followed by indicated compound treatment for an additional 6 h. β-Actin was used as the loading control. **D.** Predicted binding modes of ZSQ253 and HQY1428 obtained by molecular docking using the CR8-bound CDK12-DDB1 co-crystal structure (PDB: 6TD3) as a template. The crystallographic binding mode of CR8 is shown for comparison. Dashed lines indicate protein-ligand interactions. Representative derived compounds and their docking scores are listed. **E.** Metabolic stability of HQY1428 and ZSQ253 in mouse liver microsomes, with verapamil as the reference compound. The percentage of compound remaining over time and estimated half-lives are shown. **F.** Time-dependent degradation of cyclin K and CDK12 in SKOV3 cells treated with HQY1428 (0.1 μM), ZSQ253 (0.1 μM), or CR8 (0.1 μM). Protein levels were quantified by densitometric analysis of the immunoblots shown in Figure 3B. Curves were fitted to estimate the apparent depletion half-times (t1/2). **G.** Immunoblotting analysis of cyclin K in SKOV3 cells pretreated with MLN4924 (0.5 μM) for 2 h followed by treatment with the indicated concentrations of HQY1428, ZSQ253, CR8 or ZSQ2549 for 6 h in the continued presence of MLN4924. GAPDH was used as the loading control. **H.** SKOV3 cells with *DDB1*, *RBX1*, *CUL4A*, or *CUL4B* knocked out were incubated with or without HQY1428 (0.3 μM), ZSQ253 (0.3 μM), or CR8 (0.3 μM) for 6 h, and the indicated proteins were detected by immunoblotting. GAPDH was used as the loading control.

### Supplementary Figure 3.

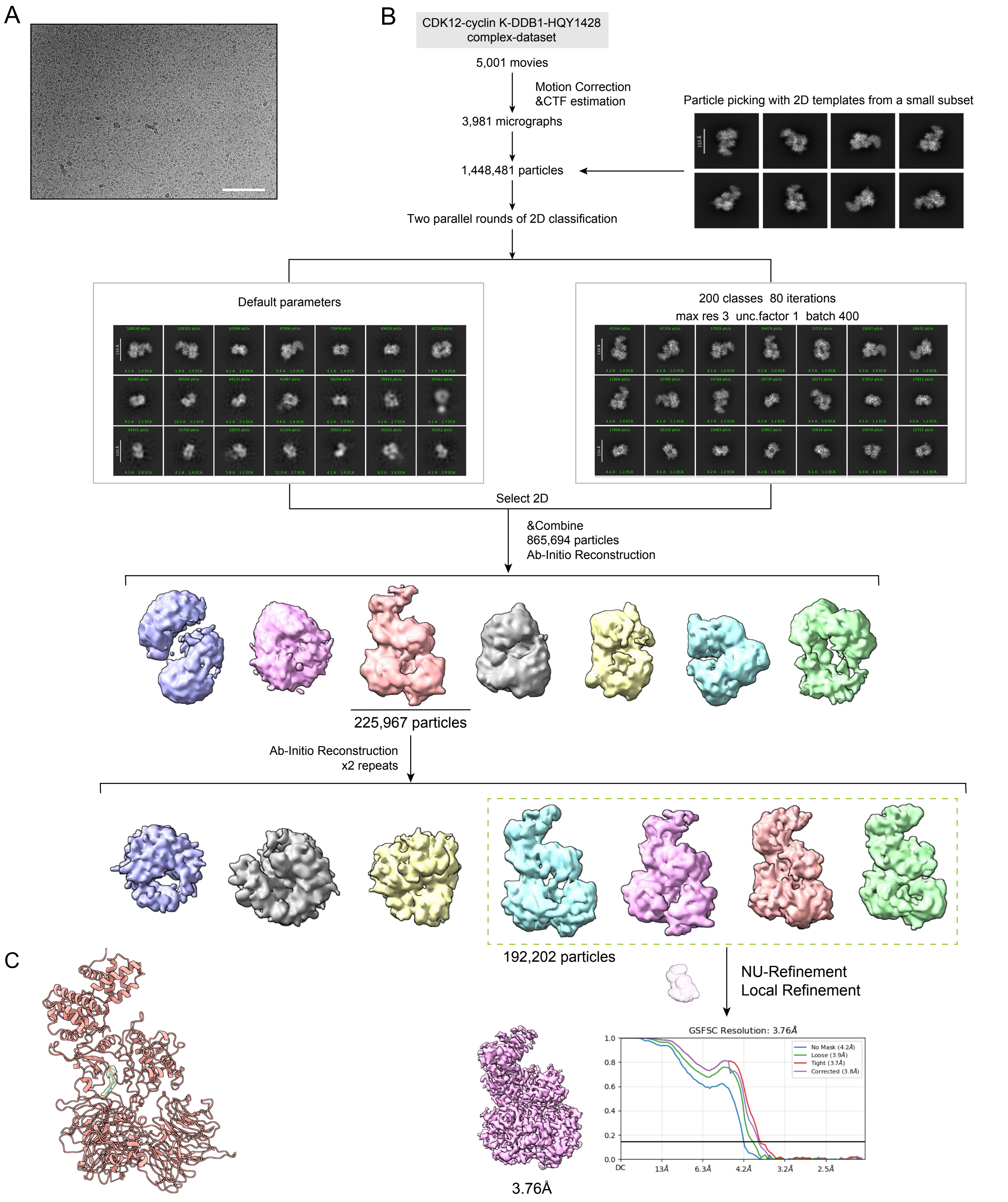


**Supplementary Figure 3.**
**A.** Representative cryo-electron microscopy (cryo-EM) micrograph of the CDK12-cyclin K-DDB1-HQY1428 complex. Scale bar, 100 nm. **B.** Image-processing workflow of CDK12-cyclin K-DDB1-HQY1428 complex. **C.** The modeled protein structure fitted to the cryo-EM density map is shown in pink, and the compound is shown in yellow.

### Supplementary Figure 4.

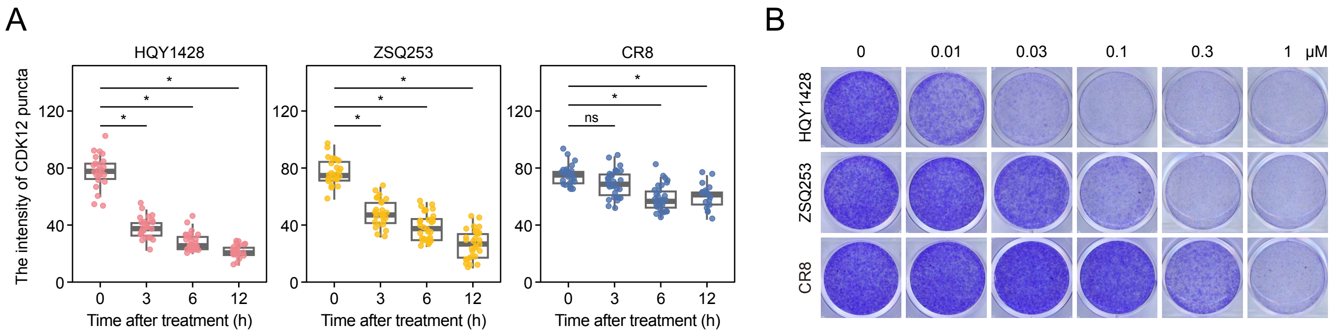


**Supplementary Figure 4.**
**A.** Quantification of CDK12 puncta intensity in SKOV3 cells treated with HQY1428 (0.1 μM), ZSQ253 (0.1 μM), or CR8 (0.1 μM) for the indicated times. Each point represents a single cell. *$P < 0.05$; ns, not significant; ANOVA followed by Tukey's post-test. **B.** SKOV3 cells were treated with HQY1428, ZSQ253, or CR8 at various concentrations for 5 days. Cell viability was determined by crystal violet staining.

### Supplementary Figure 5.

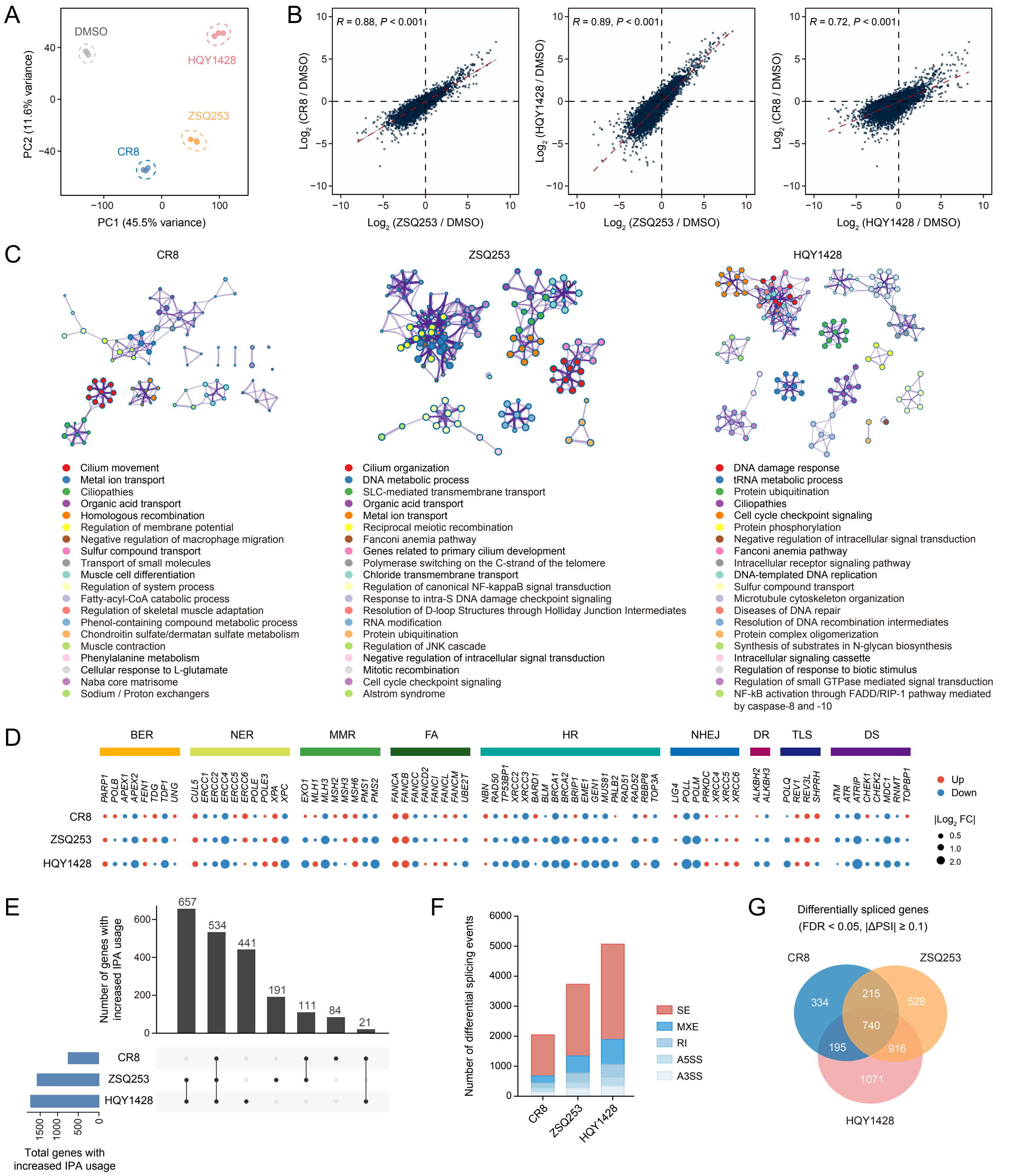


**Supplementary Figure 5.**
**A.** Principal component analysis (PCA) of RNA-seq data from SKOV3 cells treated with DMSO, CR8, ZSQ253, or HQY1428 (0.1 μM each) for 6 h. Each point represents one sample. **B.** Pairwise correlations of transcriptome-wide log2 fold changes induced by CR8, ZSQ253, or HQY1428 relative to DMSO-treated controls in SKOV3 cells. Pearson correlation coefficients (R) and P values are presented. **C.** Cytoscape network plots showing enriched terms of differentially expressed genes in SKOV3 cells treated with CR8, ZSQ253, or HQY1428 relative to DMSO. Each node represents an enriched term and is colored by cluster. Edges connect similar terms. A representative term is used to name each cluster. **D.** Bubble plot showing expression changes of core DDR genes from different DDR pathways in SKOV3 cells treated with CR8, ZSQ253, or HQY1428 relative to DMSO. Bubble size indicates absolute $\log_2$ fold change ($|\log_2 FC|$), and bubble color indicates gene upregulation (red) or downregulation (blue). BER, base excision repair; NER, nucleotide excision repair; MMR, mismatch repair; FA, Fanconi anemia; HR, homology-dependent recombination; NHEJ, non-homologous end joining; DR, direct repair; TLS, translesion synthesis; DS, damage sensors. **E.** UpSet plot showing the numbers and overlap of genes with significantly increased IPA usage in SKOV3 cells treated with CR8, ZSQ253, or HQY1428 relative to DMSO-treated controls. Horizontal bars indicate the total number of genes identified in each treatment, and vertical bars indicate the number of genes unique to or shared among the indicated treatments. **F.** Bar graph showing the number of differential alternative splicing events (FDR < 0.05, |ΔPSI| ≥ 0.1) in SKOV3 cells treated with CR8, ZSQ253, or HQY1428 relative to DMSO. PSI, percent spliced in; A3SS, alternative 3' splice site; A5SS, alternative 5' splice site; MXE, mutually exclusive exons; RI, retained intron; SE, skipped exon. **G.** Venn diagram showing the overlap of differentially spliced genes in SKOV3 cells treated with CR8, ZSQ253, or HQY1428.

### Supplementary Figure 6.

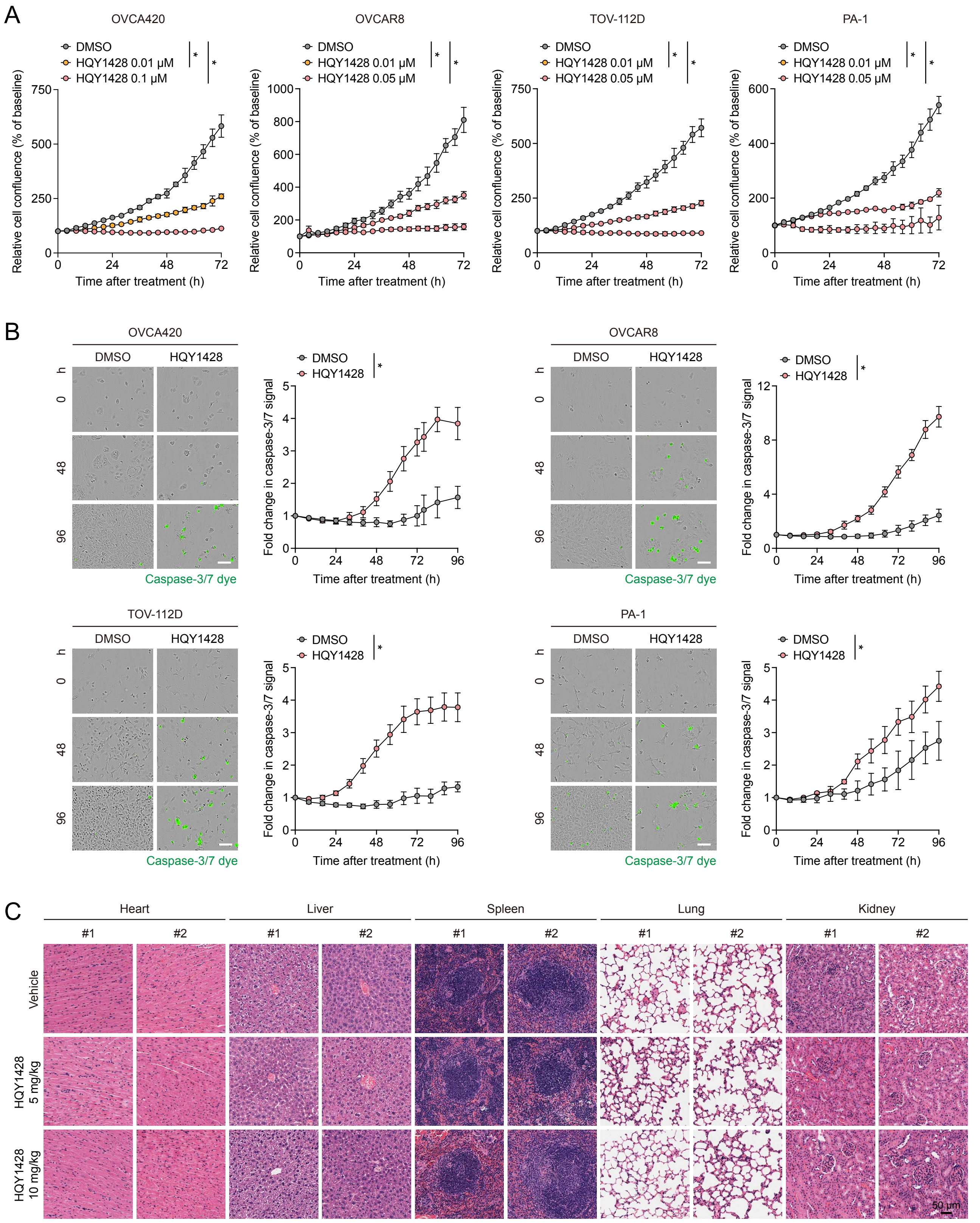


**Supplementary Figure 6.**
**A.** OVCA420, OVCAR8, TOV-112D, and PA-1 cells were treated with DMSO or the indicated concentrations of HQY1428, and cell growth was monitored for 72 h using the Incucyte live-cell imaging system. Data are presented as mean ± s.d. (n = 4). **P* < 0.05, two-way ANOVA. **B.** Representative images of OVCA420, OVCAR8, TOV-112D, and PA-1 cells treated with DMSO or HQY1428 (0.1 μM for OVCA420 and 0.01 μM for the other cell lines) in the presence of caspase-3/7 green detection reagent. Changes in caspase-3/7 fluorescence over 96 h are shown on the right. Data are presented as mean ± s.d. (n = 4). **P* < 0.05, two-way ANOVA. Scale bar, 200 μm. **C.** Representative images of hematoxylin and eosin (H&E) staining of heart, liver, spleen, lung, and kidney tissues collected from mice orally treated with vehicle control (2.5% v/v DMSO and 97.5% v/v 30% SBE-β-CD) or HQY1428 (5 or 10 mg/kg/day) for 21 days.

### Supplementary Figure 7.

A

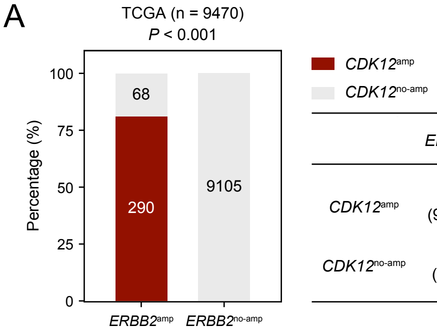


| | *ERBB2*amp | *ERBB2*no-amp |
|---|---|---|
| *CDK12*amp | 290 (97.64%) | 7 (2.36%) |
| *CDK12*no-amp | 68 (0.74%) | 9105 (99.26%) |

B

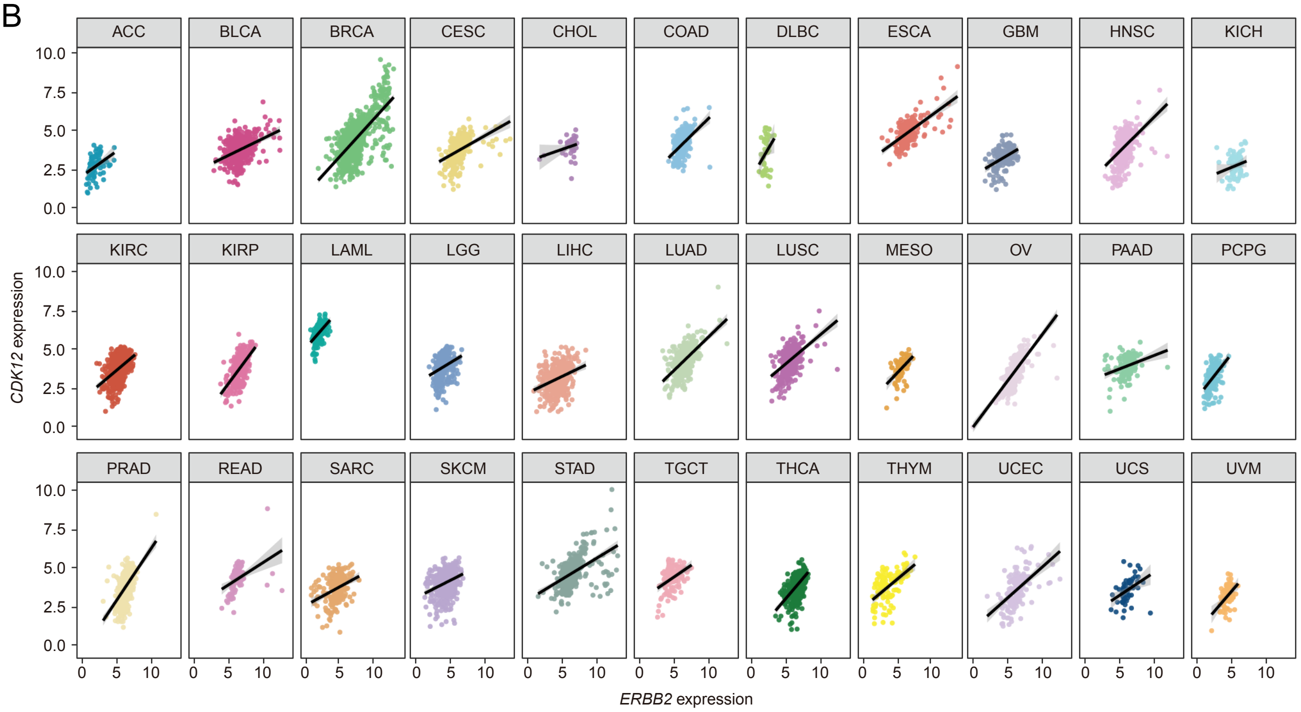


**Supplementary Figure 7.**
**A.** *CDK12* amplification status in *ERBB2*-amplified and non-amplified tumor samples from The Cancer Genome Atlas (TCGA) database (n = 9,470). Sample counts and percentages are shown in the table on the right. **B.** Correlations between *CDK12* and *ERBB2* gene expression within individual tumor types in the TCGA database.

### Supplementary Figure 8.

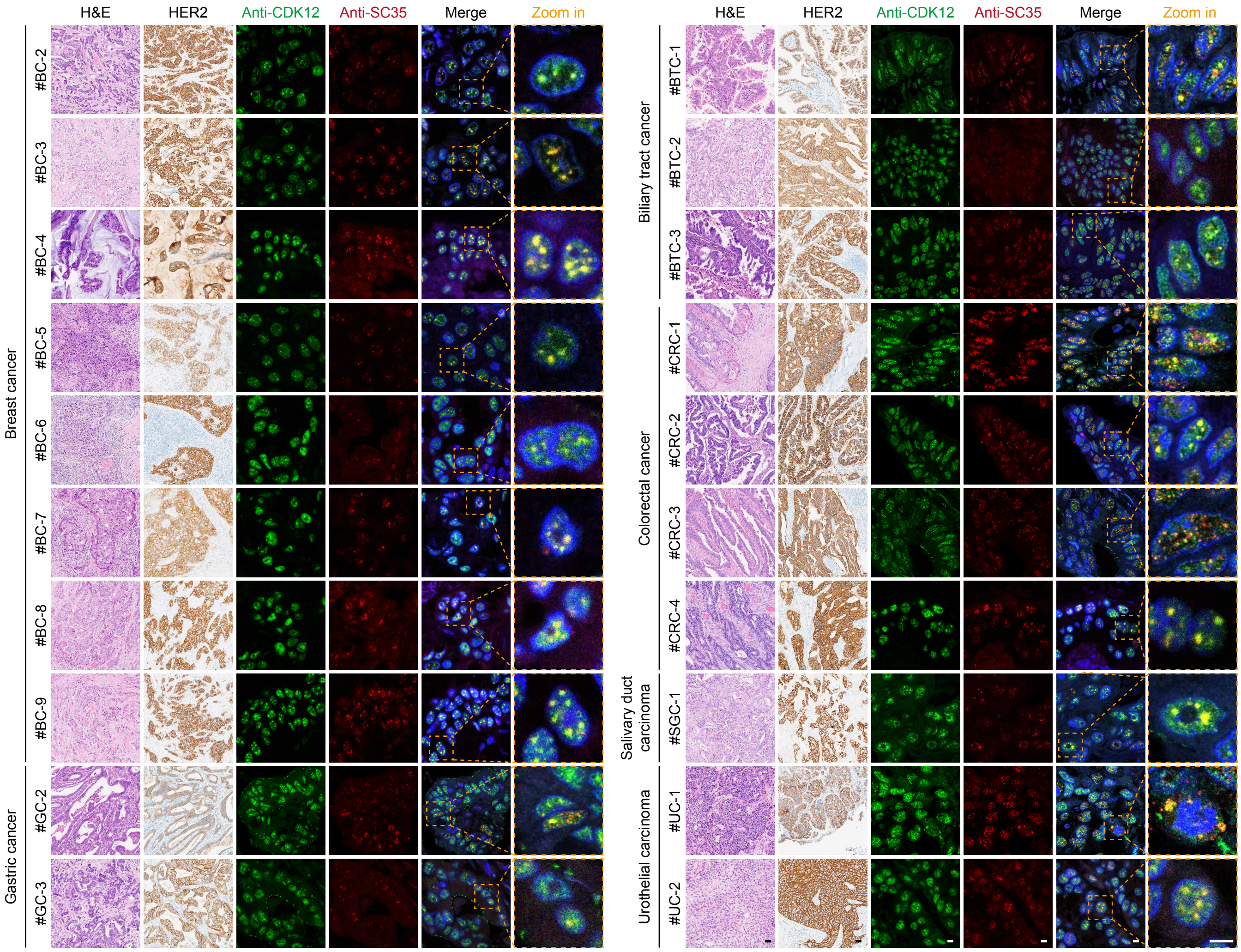


**Supplementary Figure 8.**
Representative images of hematoxylin and eosin (H&E) staining, immunohistochemistry (IHC) staining for HER2, and immunofluorescence (IF) staining for CDK12 (green) and SC35 (red) in HER2-positive breast, gastric, biliary tract, colorectal, salivary duct, and urothelial tumor sections. Cell nuclei were counterstained with DAPI (blue). Boxed regions are enlarged on the right. Scale bar, 50 μm (H&E and IHC) or 10 μm (IF).

### Supplementary Figure 9.

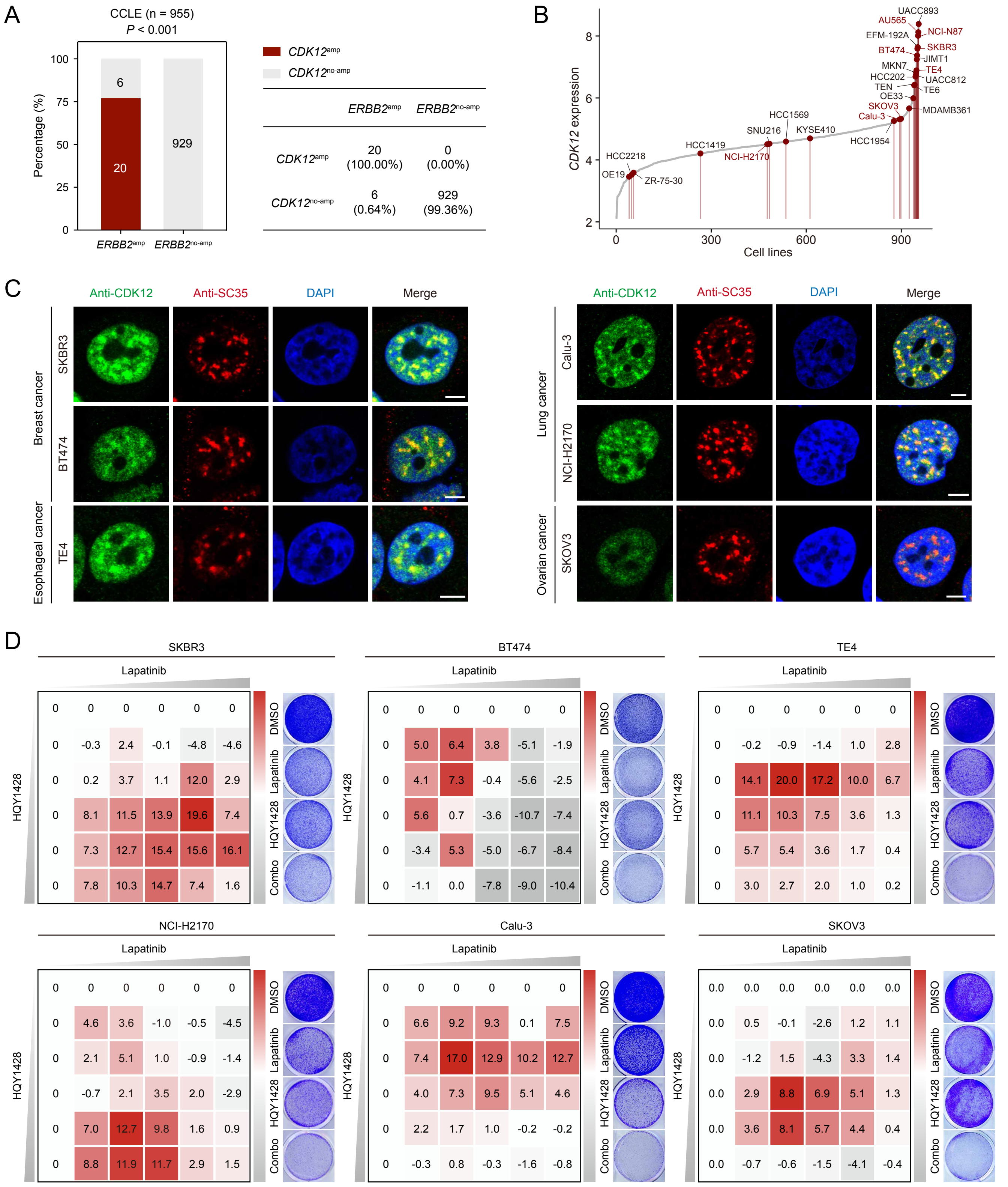


**Supplementary Figure 9.**
**A.** *CDK12* amplification status in *ERBB2*-amplified and non-amplified cancer cell lines from the Cancer Cell Line Encyclopedia (CCLE) database (n = 955). Cell line counts and percentages are shown in the table on the right. **B.** The ranking of *CDK12* gene expression across cancer cell lines in the CCLE database. Cell lines with *ERBB2* amplification are highlighted as red dots. Cell lines used in this study are labeled in red text. **C.** Representative immunofluorescence images of CDK12 (green) and SC35 (red) in breast cancer, esophageal cancer, lung cancer, and ovarian cancer cell lines with *ERBB2* amplification. Cell nuclei were counterstained with DAPI (blue). Scale bar, 5 μm. **D.** Bliss synergy score heatmaps for lapatinib and HQY1428 combinations in SKBR3, BT474, TE4, NCI-H2170, Calu-3, and SKOV3 cells. Representative crystal violet staining of cells treated with DMSO, lapatinib, HQY1428, or their combination is shown alongside each heatmap.

### Supplementary Figure 10.

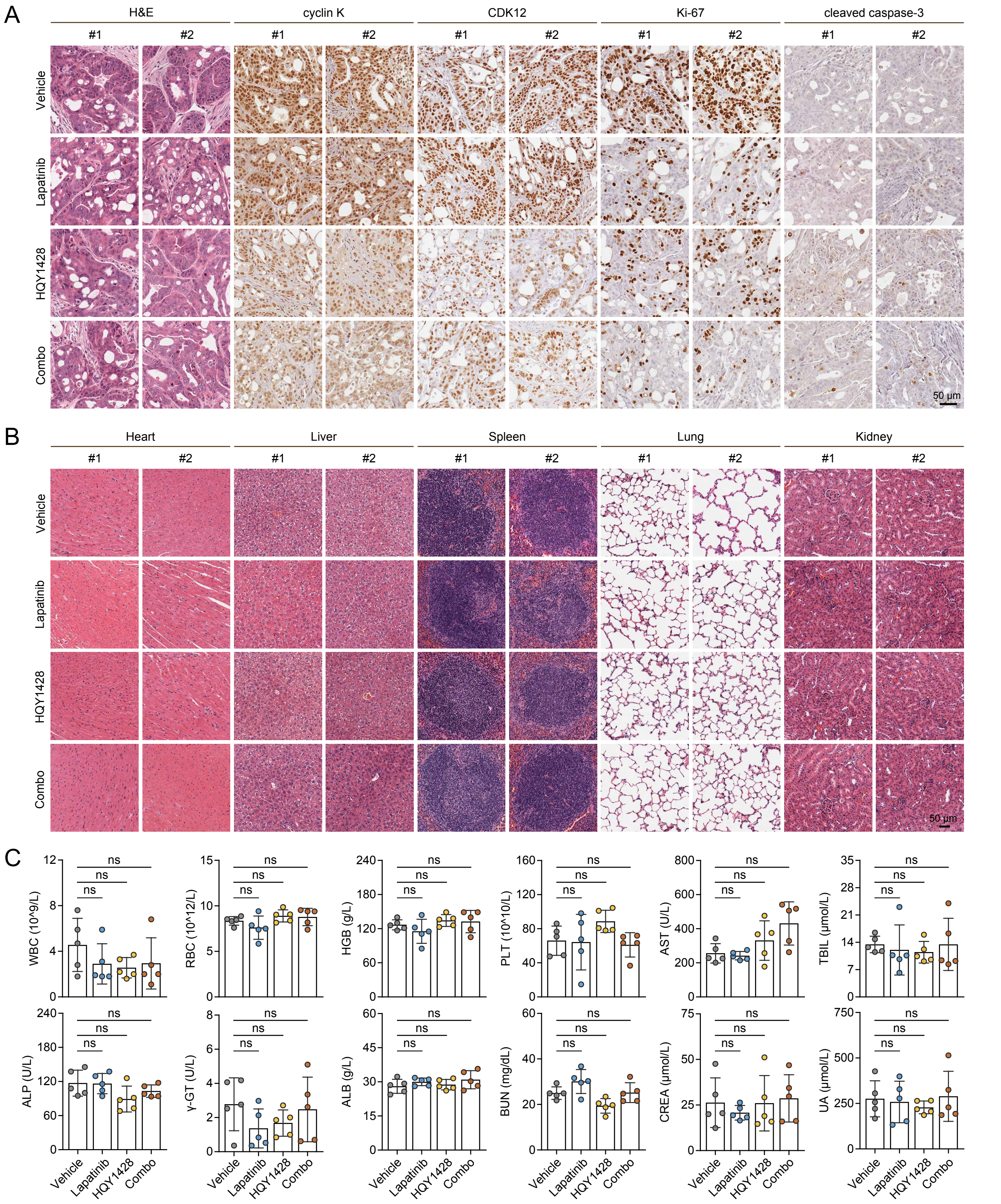


**Supplementary Figure 10.**
**A.** Representative images of hematoxylin and eosin (H&E) staining and immunohistochemistry (IHC) staining for cyclin K, CDK12, Ki-67, and cleaved caspase-3 in NCI-N87 xenograft sections from mice treated with vehicle control, lapatinib, HQY1428, or their combination. Scale bar, 50 μm. **B.** Representative images of H&E staining of heart, liver, spleen, lung, and kidney tissues collected from mice orally treated with vehicle control (0.5% hydroxypropyl methylcellulose and 0.1% Tween 80), lapatinib (30 mg/kg/day), HQY1428 (10 mg/kg/day), or their combination. Scale bar, 50 μm. **C.** Mice were orally treated with vehicle control, lapatinib, HQY1428, or their combination, followed by blood tests measuring white blood cell count (WBC), red blood cell count (RBC), hemoglobin (HGB), platelet count (PLT), aspartate aminotransferase (AST), total bilirubin (TBIL), alkaline phosphatase (ALP), gamma-glutamyl transferase (γ-GT), albumin (ALB), blood urea nitrogen (BUN), creatinine (CREA), and uric acid (UA). Data are presented as mean ± s.d. (n = 5). ns, not significant, ANOVA followed by Tukey's post-test.
